# A novel phylogenomics pipeline reveals extensive topological conflict in the evolution of the angiosperm order Cucurbitales

**DOI:** 10.1101/2023.10.27.564367

**Authors:** Edgardo M. Ortiz, Alina Höwener, Gentaro Shigita, Mustafa Raza, Olivier Maurin, Alexandre Zuntini, Félix Forest, William J. Baker, Hanno Schaefer

## Abstract

High-throughput sequencing data, such as target capture, RNA-Seq, genome skimming, and high-depth whole genome sequencing, are used for phylogenomic analyses. Integrating these mixed data types into a single phylogenomic dataset requires several bioinformatic tools and significant computational resources. Here, we present CAPTUS, a novel pipeline to analyze mixed data efficiently. CAPTUS assembles these data types, searches for loci of interest, and produces paralog-filtered alignments. If reference target loci are not available for the studied taxon, CAPTUS can also be used to discover new putative homologs via sequence clustering. Compared to other software, CAPTUS allows the recovery of a greater number of more complete loci across more species. We apply CAPTUS to assemble a comprehensive dataset, comprising the four types of sequencing data for the angiosperm order Cucurbitales, a clade of about 3,100 species in eight mainly tropical plant families, including begonias (Begoniaceae) and gourds (Cucurbitaceae). Our phylogenomic results support the currently accepted circumscription of Cucurbitales except for the position of the holoparasitic Apodanthaceae, which group with Rafflesiaceae in Malpighiales. A subset of mitochondrial gene regions supports the earlier divergence of Apodanthaceae in Cucurbitales. However, the nuclear regions and majority of mitochondrial regions place Apodanthaceae in Malpighiales. Within Cucurbitaceae, we confirm the monophyly of all currently accepted tribes but also reveal hybridization and incomplete lineage sorting both in Cucurbitales and within Cucurbitaceae. We show that contradicting results among earlier phylogenetic studies in Cucurbitales can be reconciled when accounting for gene tree conflict and demonstrate the efficiency of CAPTUS for complex datasets.

---

Modern sequencing technology allows the rapid accumulation of large amounts of genomic data at low cost. The analysis of these raw data, however, can still be a bottleneck. Researchers who cannot afford to pay for a bioinformatics expert rely on user-friendly analysis pipelines that are accessible without specialist training. As a consequence, these pipelines are of enormous importance for the field of phylogenomics. Among them, we find PHYLUCE (Faircloth 2016) and UCEasy (Ribeiro et al. 2021) specialized in the analysis of Ultra Conserved Elements (UCEs), HYBPIPER (Johnson et al. 2016) and SECAPR (Andermann et al. 2018) aimed at target capture data, PhyloHerb (Cai et al. 2022) for genome skimming data or more generalist pipelines, such as PhyloTol (Cerón-Romero et al. 2019), which analyzes previously assembled transcriptomes or annotated genomes and attempts to find all the possible gene families in a taxonomic group.

HYBPIPER and SECAPR are the most commonly used pipelines for target capture data in plant phylogenomics, however, they are designed to process one sample at a time, only making use of multiple computer cores at certain stages [e.g., when assembling sequences *de novo* with SPADES (Bankevich et al. 2012)]. To process multiple samples simultaneously, the users must rely on external tools such as GNU PARALLEL (Tange 2021) or additional containerization tools (Jackson et al. 2023). This lack of native parallelization capabilities can lead to suboptimal utilization of the computing resources in high-performance clusters or workstations and reduce accessibility for non-experts, potentially causing repeatability issues due to command inconsistencies across samples or human error. Also, since these pipelines were optimized for target capture data, their processing times for other data types can be exceptionally long. In contrast, most specialized user-friendly analysis pipelines designed for restriction site associated DNA (RAD) data, are aimed to efficiently process many samples in parallel, e.g., IPYRAD (Eaton and Overcast 2020), DDOCENT (Puritz et al. 2014), and to a lesser extent, Stacks (Rochette et al. 2019). For example, IPYRAD (Eaton and Overcast 2020) uses a single or just a few consistent commands, that, combined with relatively short processing times, allow for rapid testing of alternative settings. IPYRAD allows users to summarize results of each processing step so that they can decide on the most appropriate settings needed for following steps.

With these features in mind, we developed CAPTUS, a pipeline written entirely in Python and aimed at building phylogenomic datasets from multiple types of high-throughput sequencing data (target capture, RNA-Seq, genome skimming, and high-depth whole genome sequencing). CAPTUS makes extensive use of Python’s native parallel computing to process many samples simultaneously in a consistent manner. This parallelization can be automated or configured manually according to user needs. Each step has a simple basic command syntax (although it can be customized with many options), provides extensive reports in HTML to guide the settings of the next step, and is fully logged for repeatability. CAPTUS is able to extract nuclear, mitochondrial, and plastid proteins considering the different genetic codes and gene characteristics of each genomic compartment, as well as any other type of DNA targets (e.g., ribosomal RNA, introns, intergenic spacers) in a single command. In HYBPIPER, this would require as many runs as target types; moreover, SECAPR is not able to perform translated searches on the assemblies, limiting its targets to nucleotide sequences only. During the alignment stage, CAPTUS can also filter paralogs by taking advantage of reference target files that contain multiple sequences per locus such as the Angiosperms353 target file, a set of genes targeted by a universal probe set (Johnson et al. 2019), and its more taxonomically comprehensive version Mega353 (McLay et al. 2021). The final outputs of CAPTUS are multiple sequence alignments (MSAs) for each reference locus found in the samples. The MSAs are organized by genomic compartment and format (amino acid, nucleotide, etc.), and multiple versions of each are provided (i.e., unfiltered, paralog-filtered, untrimmed, and trimmed) so the user can select which type of alignment to use for phylogenetic inference. Additionally, a special feature of CAPTUS is the search for novel conserved markers by clustering contigs that received no hits from the reference target loci across samples, thus making use of data that otherwise would be ignored.

We demonstrate the potential of CAPTUS with the example of the flowering plant order Cucurbitales, according to APG IV (The Angiosperm Phylogeny Group 2016) a clade of eight plant families with diversity centers in the Tropics: Anisophylleaceae, Apodanthaceae, Begoniaceae, Coriariaceae, Corynocarpaceae, Cucurbitaceae, Datiscaceae, and Tetramelaceae. Together, they include 110 genera with more than 3,100 species, about 2,000 of them in the mega-diverse genus *Begonia* in Begoniaceae (Goodall-Copestake et al. 2009) and 1,000 in Cucurbitaceae (Stevens 2001; Schaefer 2020). The genus *Begonia* is of great horticultural importance while Cucurbitaceae include some of the most important vegetable and fruit crops worldwide, like cucumber and watermelon (Schaefer and Renner 2011a). Morphologically, the taxa of Cucurbitales are rather diverse, ranging from endophytic holoparasites in Apodanthaceae (Bellot and Renner 2014) to annual and perennial herbs in Datiscaceae and Begoniaceae, to trees and shrubs in Anisophylleaceae, Corynocarpaceae, Coriariaceae, and Tetramelaceae, and finally to woody or herbaceous climbers and creepers in Cucurbitaceae (Schaefer and Renner 2011a; Schaefer 2020).

A number of studies addressed phylogenetic problems in Cucurbitales in the past three decades. The earliest comprehensive seed plant phylogeny based on *rbcL* sequences (Chase et al. 1993) already identified a clade uniting the genera *Coriaria* (Coriariaceae), *Datisca* (Datiscaceae), *Luffa* (Cucurbitaceae), *Cucurbita* (Cucurbitaceae), *Begonia* (Begoniaceae), and *Tetrameles* (Tetramelaceae). Zhang et al. (2006) produced the first phylogeny estimate focused on Cucurbitales based on nine plastid, nuclear and mitochondrial loci and confirmed the results of Chase et al. (1993) while adding Anisophylleaceae and Corynocarpaceae to the Cucurbitales clade. The work of Barkmann et al. (2007) suggested another surprising addition to the Cucurbitales clade: based on mitochondrial *matR* sequence data, the holoparasitic Apodanthaceae were placed in Cucurbitales. Filipowicz and Renner (2010) increased the *matR* dataset of Barkmann et al. (2007), added a nuclear 18S dataset, and confirmed that the holoparasitic Apodanthaceae were best placed in Cucurbitales as sister lineage to all other families. Schaefer and Renner (2011b) inferred phylogenetic relationships in the order based on 14 DNA regions from all three genomes, confirming the position of Apodanthaceae based on *matR*, while their nuclear 18S-ITS-26S dataset placed Apodanthaceae outside of Cucurbitales. Other studies targeted individual families: Zhang et al. (2007) provided the first comprehensive phylogeny estimate for Anisophylleaceae, Kocyan et al. (2007) and Schaefer et al. (2009) for Cucurbitaceae, Goodall-Copestake et al. (2009) for Begoniaceae, and Renner et al. (2020) for Coriariaceae. In recent years, several phylogenomic studies contributed to an even better understanding of the evolutionary relationships within the family Cucurbitaceae.

Bellot et al. (2020) analyzed entire plastomes plus a set of 57 single-copy nuclear genes and the ITS region for 29 species from all but one of the accepted tribes and detected four nodes with conflicting phylogenetic signals. With an impressive set of 136 transcriptomes and full genomes of Cucurbitaceae, representing c. 50% of the genera, Guo et al. (2020) produced a well-supported phylogeny estimate for the family that conflicted with earlier studies in several positions (Kocyan et al. 2007; Schaefer et al. 2009; Schaefer and Renner 2011b). Finally, our recent angiosperm-wide analysis (Zuntini et al. 2024) in the framework of the Plant and Fungal Trees of Life (PAFTOL) project (Baker et al. 2022) with an almost complete representation of Cucurbitales at the genus level placed the holoparasitic Apodanthaceae outside Cucurbitales in Malpighiales and found a sister relationship between Cucurbitales and Rosales, challenging the results of most earlier studies.

In this study, we provide a detailed description of our new analysis pipeline CAPTUS and demonstrate how it can be used not only to extract the Angiosperms353 genes (Johnson et al. 2019), but also to derive a new set of thousands of nuclear genes from transcriptomic data and to extract all mitochondrial genes and entire plastomes for the Cucurbitales. The resulting phylogenomic datasets were analyzed with coalescent and concatenation methods to infer a complete genus-level nuclear phylogeny as well as plastid and mitochondrial phylogenies and test the earlier results for the topology of the order. We compare the performance and efficiency of CAPTUS against the most widely used pipeline in plant phylogenomics, HYBPIPER, which is also able to use amino acid sequences as reference targets and can handle multiple reference target sequences per locus.

## Materials & Methods

### Sampling

We used a total of 125 Cucurbitales samples. Of these, 113 are target capture sequencing samples produced for this study, but have been published previously in broader angiosperm phylogenetic analyses (Baker et al. 2022; Zuntini et al. 2024), while five target capture samples and seven high-depth whole genome sequencing samples have not been previously published (Table S1). Additionally, we downloaded sequence data for 327 samples (Cucurbitales and outgroups) available in NCBI’s SRA: 240 RNA-Seq samples, 48 genome skimming samples (<30x), 31 high-depth whole genome sequencing samples (>30x), and eight additional target capture samples (Table S2). The entire dataset comprises 342 samples of Cucurbitales sensu APG IV, representing all 110 currently accepted genera and 249 of the c. 3,100 species. To test the results of our recent angiosperm-wide analysis (Zuntini et al. 2024) and the study of Alzate et al. (2024), who found Apodanthaceae outside Cucurbitales as sister of Malpighiales or nested in Malpighiales, we selected a taxonomically broad outgroup that includes 110 samples covering taxa across the three other orders of the nitrogen-fixing clade (25 species in eight families of Rosales, 11 species in six families of Fagales, and 28 species in three families of Fabales), as well as 31 species in 23 families of Malpighiales, six species in six families of Malvales, three species of Huaceae (Huales), and three species of Vitaceae (Vitales).

### DNA Extraction and Sample Preparation

DNA samples were taken from the DNA banks of the Plant Biodiversity group at the Technical University of Munich and the Royal Botanic Gardens, Kew. Tissue samples for DNA extractions came from the herbaria of the Royal Botanic Gardens, Kew (K), Muséum National d’Histoire Naturelle in Paris (P), Botanische Staatssammlung München (M), and the Technical University of Munich (TUM).

Several different methods were used for DNA extraction from herbarium material intended for target capture: (i) a CTAB-chloroform-based (Doyle and Doyle 1987) protocol with ethanol washes and a cesium chloride/ethidium bromide density gradient cleaning and dialysis, (ii) a modified CTAB protocol (Doyle and Doyle 1987) followed by an Agencourt AMPure XP bead clean-up (Beckman Coulter, Indianapolis, Indiana, USA), (iii) an SDS-based protocol using Magen HiPure SF Plant DNA Kit (Angen Biotech Co., Ltd, Guangzhou, China) using magnetic beads for extraction and cleaning, and (iv) the manufacturers protocol of the CTAB-based NucleoSpin Plant II Extraction Kit (MACHEREY-NAGEL GmbH & Co. KG, Düren, Germany) using silica columns for binding the DNA. Depending on the available herbarium material, we used 20-160 mg of dry material for extraction.

To measure DNA quality, we evaluated fragment size distribution using an Agilent Technologies 4200 TapeStation System with Genomic DNA ScreenTapes (Agilent Technologies, Santa Clara, California, USA) or by electrophoresis in 1% agarose gel. DNA was quantified using a Quantus™ Fluorometer (Promega Corporation, Madison, WI, USA) or with a Qubit 3.0 Fluorometer (Invitrogen, Walthan, Massachusetts, USA).

The seven samples intended for high-depth whole genome sequencing were extracted with one of the following methods: (i) the standard protocol of the CTAB-based NucleoSpin Plant II Extraction Kit (MACHEREY-NAGEL GmbH & Co. KG, Düren, Germany), and (ii) the standard protocol of the MagBind® Plant DNA plus 96 Kit (Omega Bio-tek Inc.

Norcross, Georgia, USA). These seven DNA extractions were sent to GENEWIZ Germany GmbH (Leipzig, Germany), where libraries were prepared with the NEBNext Ultra II Library Prep Kit for Illumina (New England BioLabs, Ipswich, Massachusetts, USA) and sequenced on a NovaSeq 6000 platform (Illumina, San Diego, California, USA) with a paired-end 2x150 configuration.

### Library Preparation, Pooling, Target Enrichment, and Sequencing

Shearing was only performed for samples with fragment sizes above 350 bp using a Covaris M220 Focused-ultrasonicator with Covaris microTUBES AFA Fiber Pre-Slit Snap-Cap (Covaris, Woburn, Massachusetts, USA). Libraries were prepared with the DNA NEBNext UltraTM II Library Prep Kit, including size selection of DNA fragments with a length of 300 to 400 bp using SPRI beads (Agencourt AMPure XP Bead Clean-up; Beckman Coulter, Indianapolis, IN, USA.

Libraries were grouped and pooled by similar DNA quality and quantity. For target enrichment via hybridization of the pooled libraries, we used the Arbor Biosciences myBaits Target Capture Kit, “Angiosperms353 v1” (Catalog #3081XX), following the manufacturer’s manual Version 4.01 (myBaits® Kit Manual – Arbor Biosciences 2020). Hybridized libraries were PCR-amplified using the NEBNext Ultra II Q5 Master Mix (New England BioLabs, Ipswich, Massachusetts, USA). Quality and quantity of the library pools were evaluated with TapeStation and Qubit respectively as above, and then normalized and pooled for sequencing of paired-end 2x300 reads on a MiSeq platform (Illumina, San Diego, California, USA) at the Jodrell Laboratory, Royal Botanic Gardens, Kew or paired-end 2x150 reads at Macrogen Europe B.V. in Amsterdam (Macrogen, Inc., Seoul, Korea) with a HiSeq system (Illumina, San Diego, California, USA).

### Sequence Analysis Workflow

CAPTUS, implemented in Python 3, is freely available and maintained through GitHub (https://github.com/edgardomortiz/Captus) including a full documentation (https://edgardomortiz.github.io/captus.docs/). It can be installed directly from Bioconda (https://bioconda.github.io/recipes/captus/README.html). The workflow of CAPTUS consists of four steps controlled by their respective modules called `clean`, `assemble`, `extract`, and `align`, which are typically run in that order (Fig. 1).

**FIGURE 1.**
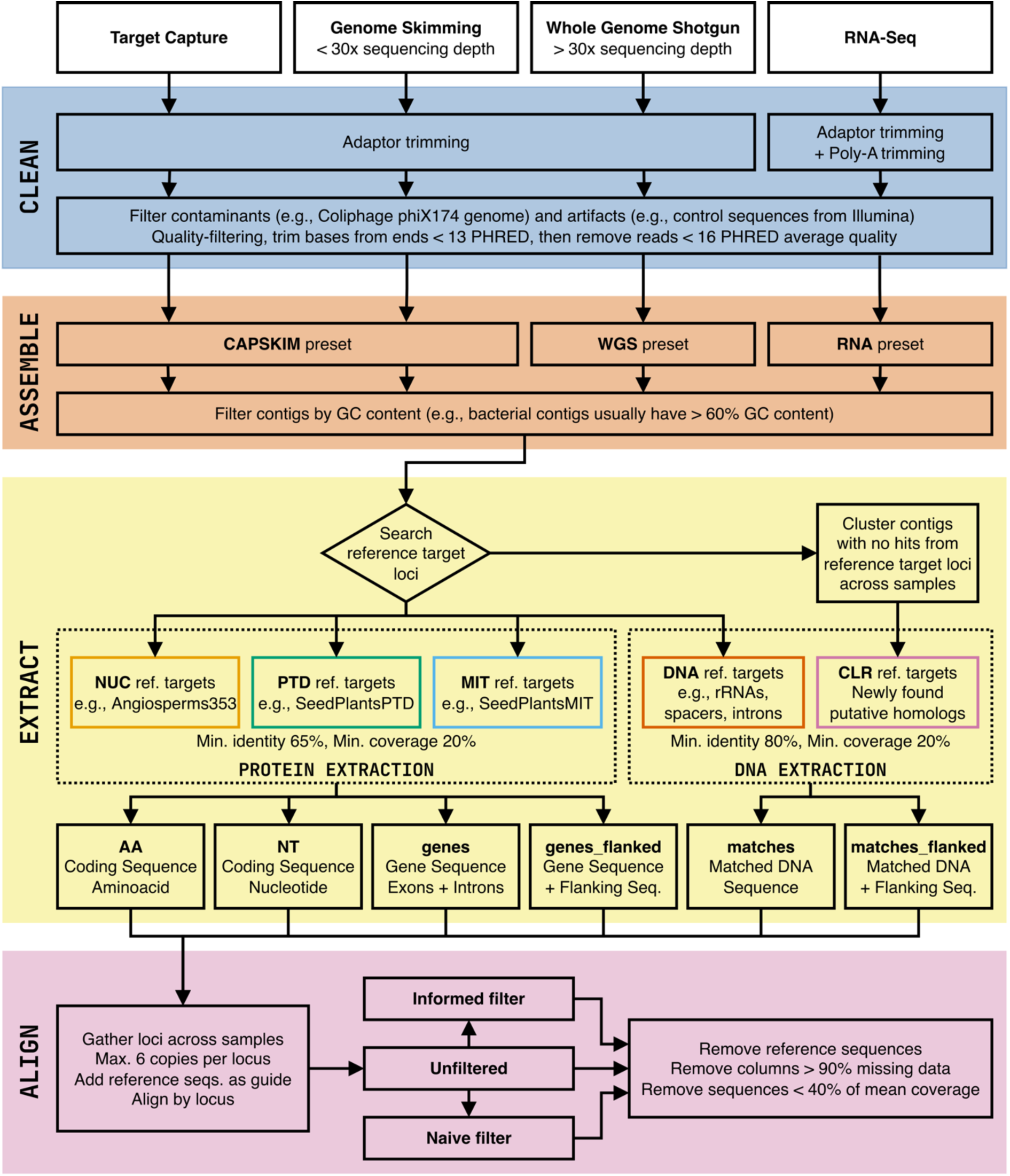
Workflow of the analysis pipeline CAPTUS. The default thresholds shown can be changed by the user via command options. NUC: nuclear proteins, PTD: plastidial proteins, MIT: mitochondrial proteins, DNA: miscellaneous DNA targets, CLR: DNA targets found via clustering Alternatively, the analysis can be started at different points depending on the sample data provided. If raw reads are provided one should start with the `clean` module, if the reads have already been cleaned, they can be directly analyzed with the `assemble` module, and if previously assembled data or reference genomes are provided the analysis can start using the `extract` module. CAPTUS provides flexibility in combining samples that enter the workflow at different stages. For example, a phylogenomic dataset could be composed of samples represented by target capture raw sequencing reads, samples with clean RNA-Seq reads, and genomic assemblies downloaded from GenBank to increase taxon sampling.

*The* `clean` *module.---* CAPTUS uses BBDUK from BBTOOLS (Bushnell 2022) to clean raw reads. Cleaning is performed in two steps. First, adaptors and poly-A tails (when cleaning RNA-Seq reads) are trimmed in two consecutive rounds. Then, adaptor-free reads are quality-trimmed, and reads matching the phiX viral genome or known sequencing artifacts are removed using the databases provided by BBDUK. In this step, leading and trailing bases with PHRED scores <13 are trimmed and then trimmed reads with average PHRED score <16 are removed, both thresholds can be altered by the user.

When cleaning is completed, FASTQC (Andrews 2019) or its faster implementation FALCO (de Sena Brandine and Smith 2021) is run both on the raw and the cleaned reads to evaluate and compile quality statistics. Finally, CAPTUS summarizes the cleaning statistics of all the samples before and after cleaning in a single HTML report that allows a quick overview of all quality measurements.

For the Cucurbitales analysis, default settings were used for all samples in the clean module except RNA-Seq data, for which the option --rna was added to trim poly-A tails.

*The* `assemble` *module.---* The clean reads are passed to the assembly module which uses MEGAHIT for *de novo* assembly. This assembler is capable of handling very large datasets in a time-efficient manner and with greatly reduced memory requirements because of the use of a compressed representation of *de Bruijn* graphs and very efficient parallel computing algorithms. Additionally, MEGAHIT implements the concept of *mercy-kmers*, a unique strategy that allows removing spurious low-depth contigs while keeping authentic ones (Li et al. 2015, 2016). More importantly, the ability of assembling all reads simultaneously has been found to reduce the number of chimeric contigs (García-López et al. 2015).Thanks to these features, MEGAHIT is not only exceptional at assembling metagenomic datasets (its original purpose) but also single-genome datasets, and is able to produce accurate assemblies even at low sequencing depths as shown in our previous tests with small datasets (Raza et al. 2023).

We tested several combinations of MEGAHIT settings in order to optimize the assembly of other types of data. These are provided in CAPTUS under three presets: (i) `CAPSKIM` for target sequence capture, genome skimming data, or a combination of both, (ii) `RNA` for RNA-Seq data, and (iii) `WGS` for high-depth whole genome sequencing data, which includes the recently defined “Deep genome skimming” (Liu et al. 2021) (https://edgardomortiz.github.io/captus.docs/assembly/assemble/options/#--preset).

The assemble module also provides an option to subsample a fixed number of reads per sample prior to assembly, which is useful when the sequencing depth is too high for a particular sample or for speeding up assembly during trial runs.

Once assembly is completed, CAPTUS can also filter contigs based on their GC content. For example, a maximum of 60% GC would be appropriate to remove bacterial contamination in most eukaryotic genomes (Fierst and Murdock 2017). Finally, CAPTUS computes assembly statistics and produces a single HTML report for all processed samples.

Target capture and genome skimming samples of the Cucurbitales were run with the default assembly preset (--preset CAPSKIM), RNA-Seq samples were assembled using - -preset RNA, and high-depth whole genome sequencing samples using --preset WGS. To remove potential bacterial contamination, we used the option --max_contig_gc 60 for all assemblies.

*The `extract` module.---* Once assemblies are completed, they can be searched for particular loci of interest (i.e., reference target loci). CAPTUS allows the simultaneous search and extraction of five types of markers (i) nuclear proteins (`NUC`), (ii) plastid proteins (`PTD`), (iii) mitochondrial proteins (`MIT`), (iv) miscellaneous DNA regions (`DNA`), and (v) new clustering-derived putative homologs (`CLR`).

Protein extractions (`NUC`, `PTD`, and `MIT` markers) use reference target loci provided as amino acid sequences or coding sequences in nucleotides (CDS) and a different genetic code can be specified for each separately. CAPTUS uses the program SCIPIO to perform protein extractions. SCIPIO is able to automatically correct frameshifts that could result from sequencing or assembly error and it can recover proteins and their putative paralogs that are spread across several contigs by analyzing splice patterns, checking for compatible exonic boundaries and overlaps, and for consistency in the similarity of the partial hits to the target protein to reconstruct coherent gene models (Keller et al. 2008; Hatje et al. 2011). This is important, given that loci assemblies resulting from target capture data can potentially be highly fragmented, particularly when the captured target proteins contain exons that are separated by long introns. CAPTUS comes bundled with four reference target sets, two `NUC` references [curated versions of the Angiosperms353 (Johnson et al. 2019) target file and the taxonomically expanded Mega353 (McLay et al. 2021) target file], as well as a plastid `PTD` reference (SeedPlantsPTD), and a mitochondrial `MIT` reference (SeedPlantsMIT).

Our curation of the Angiosperms353 reference target file consisted in clustering the sequences at 98% identity to remove highly similar sequences and duplicates while the curation of the Mega353 reference target file consisted in removing contaminated sequences (e.g., fungal, algal, organellar sequences) and then clustering at 78% identity to reduce sequence redundancy. Thus, CAPTUS still takes advantage of the expanded taxonomic representation of the Mega353 reference targets (McLay et al. 2021) while bypassing the additional step of filtering by taxon suggested in the instructions (https://github.com/chrisjackson-pellicle/NewTargets). The `PTD` and `MIT` target references represent carefully curated sets of 83 proteins (represented by 2,194 sequences) and 45 proteins (represented by 167 sequences) respectively, found in most organellar genomes of seed plants originally downloaded from GenBank. CAPTUS also allows the utilization of BUSCO lineage databases (e.g. https://busco-data.ezlab.org/v5/data/lineages/) as target references and new target sets for organism groups other than seed plants are possible and being developed.

Miscellaneous DNA regions (`DNA`) extractions are aimed at recovering other types of `DNA` regions (e.g., complete genes with introns, non-coding regions, ribosomal genes, individual exons, RAD markers, etc.). In this case, CAPTUS uses the program BLAT (Kent 2002) to search the assembly and the hits that are found across contigs are greedily assembled and concatenated using our own code.

When a protein or DNA target is spread among several contigs, CAPTUS concatenates the contigs to reconstruct the entire locus. This procedure is carefully done as it considers the sequence identity of each fragment to the target and their mutual overlap to minimize the possibility of chimeras. If the targets are expected to be fully contained in a single contig (e.g., when using pre-assembled genomes as input) or if chimeric contigs are of high concern, contig concatenation can be disabled with `--disable_stitching`.

Additionally, CAPTUS can be used to find new putative homologs by clustering the contigs that had no hits from the reference target loci across samples. In this case, the program MMSEQS2 (Steinegger and Söding 2018) is used, and several of its settings are available through CAPTUS. Once the sequence clusters have been found, they are filtered by the number of samples in the cluster and sequence length, and representative sequences per cluster are selected as new reference targets to perform another extraction of the miscellaneous `DNA` type in CAPTUS. The output from this second round of extraction is indicated by the prefix `CLR`.

The user can specify the minimum sequence identity and coverage to perform the search of target loci within the assemblies. If several matches are found for a single target locus, they are ranked from most to least similar and complete as compared to the reference target sequence. Therefore, the best hit is recognized in the output files by having the suffix `_00` while the rest of the copies (i.e., putative paralogs) are ranked with increasing numbers (e.g., `_01`, `_02`, etc.).

The output files from protein extractions (`NUC`, `PTD`, `MIT`) are provided in four possible formats (Fig. S1a): the protein sequence in amino acids (`AA`), the coding sequence in nucleotides or CDS (`NT`), the complete gene sequence including introns (`genes`), and the complete gene sequence flanked by a fixed number of nucleotides (`genes_flanked`).

Similarly, the output from a miscellaneous DNA extraction and clustering-derived markers (`DNA`, `CLR`), have two possible formats (Fig. S1b): the segment of sequence that was matched to the reference (`matches`), and the matched segments flanked by a fixed number of nucleotides (`matches_flanked`).

Once extractions are finished, CAPTUS collects extraction statistics (e.g., recovered marker length, similarity to the reference sequence, number of paralogs, number of contigs used in the gene assembly, etc.) from all the processed samples and produces a single HTML report that provides a quick overview of these recovery indicators across loci and samples as an interactive heatmap. For example, the user can choose to plot identity to target, recovered percentage, number of copies detected, number of contigs used in the assembly of the locus and the mean depth of coverage of those contigs, among others.

For our Cucurbitales dataset, the minimum recovery percentage was set at 20%, the default in CAPTUS. We used the option `-n Angiosperms353` to extract nuclear proteins using the bundled Angiosperms353 reference targets and `-n Mega353` to extract nuclear proteins using our curated version of the taxonomically expanded Mega353. Additionally, organellar proteins were extracted with options `-p SeedPlantsPTD` and `-m SeedPlantsMIT`. Finally, we created a custom miscellaneous DNA reference by segmenting the Cucurbitales plastomes available in GenBank in 38 pieces ranging from ∼3 kbp to ∼5 kbp which we stored in a FASTA file (`Plastome38.fasta`, Appendix S1) and extracted in CAPTUS using the option `-d`.

Additionally, we created a set of 5,435 reference genes derived from publicly available transcriptomic data. We took the 240 transcriptomic assemblies from CAPTUS and removed the transcripts that had hits to organellar proteins, thereby retaining only putatively nuclear transcripts. We ran CODAN (Nachtigall et al. 2021) on the nuclear transcripts in order to find coding regions within them, searching both strands and using the `PLANTS_full` model which only emits a CDS when a complete protein, from start to stop codon, is identified within the transcript. Then we clustered the CDS across samples using the `extract` module of CAPTUS, with the following options:

> `-c --mmseqs2_method easy-cluster --cluster_mode 2 --cl_min_identity 70 --cl_seq_id_mode 1 --cl_min_coverage 66 --0cl_rep_min_len 540 --cl_min_samples 144 --cl_max_copies 4.`

These CAPTUS options retained 5,435 clusters that were at least 540 bp in length, grouped at least 144 transcriptomic samples (60%), and contained at most an average of four gene copies.

After removing within-cluster redundant sequences, the resulting reference contained 13,492 sequences representing these 5,435 CDS clusters. This reference is called from this point onwards RNA5435 (`RNA5435.fasta`, Appendix S2). The new reference targets were used in the `extract` module of CAPTUS across all 452 samples using the option `-n` RNA5435.fasta --nuc_min_score 0.15 --nuc_min_identity 70 to match the clustering identity threshold used for creating the reference.

*The* `align` *module.---* The extraction output is then processed by the alignment module. Individual markers are collected across samples and organized in separate FASTA files per locus. By default, CAPTUS collects a maximum of five paralogs per sample and per marker to be aligned. The reference sequences are also added to each locus file to serve as alignment guides in case the sequences recovered from the samples are fragmentary. Then, the alignment is performed with MAFFT (Katoh and Standley 2013) or MUSCLE5 (Edgar 2022) using default settings or by selecting one of their specific algorithms. However, if protein sequence in amino acid (`AA`) and their corresponding CDS in nucleotides (`NT`) are aligned in the same run, CAPTUS aligns the `AA` with MAFFT and then uses the `AA` alignment as template for the `NT`, thus producing a codon-aware alignment for the CDS. CAPTUS also allows the user to provide the sample(s) that should be considered as outgroup, in this case the program will place those samples as the first sequence(s) in the alignments in the order provided. This feature (`--outgroup`) takes advantage of a common feature of many phylogenetic estimation programs (e.g., IQ-TREE, RAXML, MRBAYES) which arbitrarily draw the first sample in the alignment at the root of their output trees.

Once the FASTA files are aligned, paralogs are filtered using two alternative algorithms, `naive` and `informed`. Our paralog filters are computationally efficient because they are based on sequence similarity comparisons, however their results might differ from more computationally expensive, phylogeny-informed methods of determining orthology such as ORTHOFINDER or other pipelines (Yang and Smith 2014; Emms and Kelly 2019; Morales-Briones et al. 2021). Consequently, CAPTUS also provides the alignments that include all the paralogs found per locus (`unfiltered`) so the user can use them as input for other paralog-pruning methods.

In the `naive` filter, only the best hit (i.e., most similar and complete as compared to reference target sequence, with suffix `_00` in output files) is kept for each sample and no further filtering is performed. The `informed` filtering algorithm takes advantage of reference datasets that contain multiple sequences per locus, such as the Angiosperms353 (Johnson et al. 2019) or Mega353 (McLay et al. 2021) target files as well as the ones developed for CAPTUS (SeedPlantsPTD and SeedPlantsMIT). For example, the Angiosperms353 reference target set (as well as its expansion Mega353) was derived from the 1KP Project data (Matasci et al. 2014; One Thousand Plant Transcriptomes Initiative 2019), where each gene could potentially be present in more than 1,000 species. To build a less redundant sequence collection while still covering the entire phylogenetic diversity of the angiosperms, each selected locus was clustered at ∼70% identity, usually selecting only one representative sequence per cluster for the final reference dataset. Thanks to this feature, one could expect that, when analyzing a single taxonomic group such as a family or genus, all samples would have a best match to only one sequence in the reference targets collection for a given locus (i.e., the closest relative present in the reference targets). However, a minority of samples could have a copy (paralog) that best matches a different reference sequence, producing an inconsistent paralog ranking than the rest in the studied group, and therefore the most common copy found in the group is not the chosen best hit in the minority of samples. In these cases, CAPTUS compares all recovered paralogs with the reference sequence that best matches the majority of samples and reorders the paralog ranking of the minority keeping only the copy that is most similar to that reference sequence across samples. Finally, it could also happen that a sample presents a single copy for a specific locus but with a sequence that is much more divergent than the average in the alignment (e.g., a remote paralog found in a contaminated contig). By default, CAPTUS will remove any sequence with average pairwise identity that is more than 4.0 standard deviations below the mean pairwise identity of the entire alignment, the number of standard deviations can be changed with the option `--tolerance`.

Modern species tree inference methods can deal with phylogenetic conflict caused by ILS or introgression, but are usually limited to the analysis of single-copy gene trees (Yan et al. 2022). Fortunately, in those cases, the paralog pruning method seems to be of little consequence to the final topology, where even a random selection produced results congruent with more sophisticated methods of paralog removal (Yan et al. 2022); however further tests under complex scenarios are necessary.

Moreover, recently developed coalescent methods, such as ASTRAL-PRO (Zhang et al. 2020; Zhang and Mirarab 2022), are capable of directly analyzing single-copy gene trees as well as trees of genes with paralogs (i.e., multi-copy genes), completely bypassing the need of filtering extra gene copies. To facilitate these analyses, CAPTUS not only provides the alignments with paralogs but also the file required by ASTRAL-PRO for mapping the paralog names to the names of the samples for species tree calculation, this file is unique and can be used with any of the alignment sets produced by CAPTUS.

The reference target sequences are then removed from the alignments. Finally, all the produced alignments and their filtered versions are trimmed using CLIPKIT (Steenwyk et al. 2020), which can remove alignment columns based on criteria like informativeness or proportion of missing data. By default, CAPTUS removes columns with > 90% missing data and sequences with < 40% mean coverage. Alternatively, a minimum number of sites per column instead of a percentage can be specified with `--min_data_per_column`. As in previous modules, CAPTUS computes alignment statistics as well as sample occupancy statistics from all the FASTA files along all filtering and trimming stages and produces a comprehensive HTML report that can be used to determine outlier markers or samples that should be removed or curated more carefully before proceeding to phylogenetic analysis.

For the Cucurbitales data, all extracted loci were aligned in CAPTUS using MAFFT’s most accurate algorithm E-INS-i (`--align_method mafft_genafpair`). For coding regions we used the option `-f AA,NT` to take advantage of codon-aware alignments for CDS, for Plastome38 we used `-f MA.` For trimming, we used `--min_data_per_column 6`, to keep alignment columns with six or more sequences. For the Plastome38 we also decreased the tolerance of the `informed` paralog filter to `--tolerance 2.0`.

### Phylogenetic Analyses

We chose the trimmed, codon-aligned coding sequences (format `NT`) from the extractions performed with reference targets sets Angiosperms353, Mega353 and RNA5435. For the reference targets set Plastome38 (the plastome segments) we selected the format `matches` from the CAPTUS output. We analyzed the unfiltered alignments (i.e., including paralogs) as well as the alignments resulting from the `naive` and `informed` paralog filtering strategies for nuclear markers but only the alignments filtered by the `informed` algorithm for the plastome.

*Gene tree estimation.---* For each individual nuclear gene alignment we inferred a phylogeny using IQ-TREE v. 2.2.2.6 (Minh et al. 2020b). During the run we used MODELFINDER (Kalyaanamoorthy et al. 2017) to determine the most appropriate nucleotide substitution model with `-m TEST`. Nodal support was inferred from 1,000 ultrafast bootstrap replicates with option `-bb 1000` (Hoang et al. 2018). For comparison purposes, we also estimated nuclear gene tree phylogenies using FASTTREE v. 2.1.11 (Price et al. 2010) with the manual recommended options to increase the accuracy of the search and using the GTR substitution model (`-pseudo -spr 6 -mlacc 3 -slownni -gtr`).

*Species tree estimation.---* We estimated species trees using two methods, concatenation of alignments followed by maximum likelihood estimation in IQ-TREE v. 2.2.2.6 (Minh et al. 2020b), and coalescent estimation by summarization of quartet frequencies in gene trees using ASTRAL-PRO v. 1.15.1.3 (Zhang et al. 2020; Zhang and Mirarab 2022). For concatenation, each individual locus alignment must contain a single sequence per sample (no paralogs allowed), therefore we can only apply this method to the alignments that were filtered for paralogs, while ASTRAL-PRO was applied to the filtered alignments as well as to the alignments containing multiple copies. The concatenation analyses in IQ-TREE were run with the same options as for the individual gene trees, and the loci alignments were provided as separate files in a single directory with the option `-p`, so IQ-TREE can automatically concatenate them into a supermatrix prior to analysis. The concatenation method was applied only to the set 353 nuclear genes (extracted using Angiosperm353 or Mega353), to the plastome segments (Plastome38).

For the ASTRAL-PRO analysis, the individual gene trees calculated by IQ-TREE or FASTTREE were provided as well as the required file that maps the paralog names to their corresponding sample name produced by CAPTUS (`captus-assembly_align.astral-pro.tsv`). We also increased the number of placement and subsampling rounds to 16 (`-R`), as well as the proportion of taxa subsampled (`--proportion 0.75`) and calculated alternative quartet frequencies (`-u 3`).

*Site concordance analyses.---* In order to measure concordance between alignment sites and the species tree, we performed a concordance factor analysis for each species tree (Minh et al. 2020a). The analysis was done in IQ-TREE v. 2.2.2.6 by supplying the species tree (`-t`), the folder containing the loci alignments (`-p`), and averaging the site concordance over 1,000 quartets (`--scfl 1000`).

*Phylogenetic conflict analysis of selected nodes.---* Contentious relationships in the Cucurbitales are centered around the relationships among tribes in the Cucurbitaceae as well as the relationships among Cucurbitales families. In order to visualize the amount of conflict at selected nodes in the species tree, we used the branch quartets frequencies analysis in DISCOVISTA v. 1.0 (Sayyari et al. 2018), after annotating the species belonging to each Cucurbitaceae tribe, to each of the families in Cucurbitales, and to the clades in the outgroup.

*Phylogenetic network estimation.---* We concatenated the Angiosperms353 alignments filtered by the `informed` method in CAPTUS keeping only the Cucurbitales families. This concatenated alignment was used as input for SPLITSTREE v. 4.18.2 (Huson and Bryant 2006) in order to estimate a phylogenetic network using the Neighbor Net algorithm on a matrix of uncorrected P distances.

*Mitochondrial relationships in Cucurbitales.---* The previous phylogenetic placement of the Apodanthaceae as sister to the remaining Cucurbitales was based on the signal of a *matR* dataset (Filipowicz and Renner 2010; Schaefer and Renner 2011b). To compare our contradicting nuclear phylogenomic results with a more comprehensive mitochondrial dataset, we examined the phylogenetic relationships of all 39 mitochondrial genes found in Cucurbitales. Since three genes are trans-spliced we segmented those in two or three parts for a total of 43 mitochondrial gene regions.

To obtain a full complement of mitochondrial genes while excluding potentially irrelevant distant hits we tested minimum sequence identity thresholds between 75% and 90% with CAPTUS and determined that 83% was the highest identity threshold at which all mitochondrial genes (including *matR*) could still be recovered from the Apodanthaceae samples. Additional mitochondrial genes (e.g., *rpl2*) were not recovered for Apodanthaceae above 88% minimum sequence identity.

We extracted the mitochondrial regions from our entire dataset using CAPTUS with a minimum sequence identity of 83%, minimum coverage of 30%, and allowing a maximum of two additional copies per locus (`-d SeedPlantsMIT --dna_min_identity 83 --dna_min_coverage 30 --max_paralogs 2`). The alignment was also performed in CAPTUS using MAFFT’s most thorough algorithm (`--align_method mafft_genafpair`).

The trimmed unfiltered alignments were analyzed with IQ-TREE v2.3.6 (Minh et al. 2020b) performing standard model selection by MODELFINDER, with 1,000 ultrafast bootstraps, and using a fixed seed (`-m TEST -bb 1000 -seed 142857`).

To distinguish between “host DNA” (due to contamination or perhaps horizontal gene transfer) and “parasite DNA” in the Apodanthaceae samples, we analyzed the phylogenetic affinities of the sequenced gene copies. If the different copies of all three Apodanthaceae samples grouped together and each was represented by a single copy only, they were considered “parasite DNA”. If we recovered multiple copies per gene, those that grouped with the known host plants [Fabaceae (Fabales) for *Pilostyles*; *Casearia* (Salicaceae, Malpighiales) for *Apodanthes*] were considered “host DNA” sequences while extra copies grouping elsewhere were considered “parasite DNA”. Because the curated set of target plant mitochondrial proteins in CAPTUS (`SeedPlantsMIT`) was derived exclusively from non-parasite angiosperms, most matches to “host DNA” had identities equal or above 88%.

Therefore, if an Apodanthaceae sample had a single gene copy but did not cluster with the others, we decided based on sequence similarity: sequences above 88% identity with the reference were classified as “host DNA” and below 88% as “parasite DNA”. In a few inconclusive cases the sequences were treated as “unclear”, usually when the gene tree presented very low resolution.

### Benchmarking

The locus recovery efficiency of CAPTUS on different data types was evaluated against HYBPIPER (Johnson et al. 2016) by extracting the Angiosperms353 loci (Johnson et al. 2019) from a selection of 80 samples (20 from each data type), representing all families of Cucurbitales and all tribes of Cucurbitaceae (Table S3). Since the HYBPIPER workflow does not include a read cleaning step, we used reads cleaned by CAPTUS as a common input for both pipelines. A protein target file downloaded from https://github.com/mossmatters/Angiosperms353 was used as a reference for both pipelines.

CAPTUS was run for the `assemble` module with the aforementioned presets optimized for each data type, followed by the `extract` module with default settings. HYBPIPER v. 2.1.2 was run using either BLASTX (Camacho et al. 2009) or DIAMOND (Buchfink et al. 2021), hereafter HYBPIPER-BLASTX and HYBPIPER-DIAMOND respectively, with an identity threshold of 65% (`--thresh 65`) to equalize with that of CAPTUS. For the genome skimming samples, HYBPIPER-BLASTX and HYBPIPER-DIAMOND were run with a lower coverage cutoff of 1 (`--cov_cutoff 1`), as the default cutoff of 8 was considered too stringent for this data type. The locus recovery efficiency of each pipeline was evaluated based on three metrics: running time, number of loci recovered, and total CDS length recovered. To make the running times comparable across pipelines, all commands were run on the same MacOS X system equipped with a 2.7 GHz Intel Xeon E5 processor with 24 threads and 64 GB RAM, allocating 6 threads per sample and processing 4 samples concurrently. GNU PARALLEL v. 20230122 (Tange 2021) was used to run HYBPIPER on multiple samples concurrently and to record the running time spent on each sample in a log file (`--joblog`). The number of loci recovered and the total CDS length recovered were calculated from the stats file generated by each pipeline (`captus-assembly_extract.stats.tsv` from CAPTUS and `seq_lengths.tsv` from HYBPIPER). Additionally, HYBPIPER was run for the `paralog_retriever` command with default settings to retrieve the detected paralog sequences. The paralog sensitivity of each pipeline was evaluated based on the average number of paralogs per recovered locus. To ensure a fair comparison, only paralogs exceeding 75% of the reference protein length were counted, in accordance with the default threshold of HYBPIPER. The statistical significance of the differences between pipelines was assessed using the Wilcoxon rank-sum test with Bonferroni correction.

To investigate the relationship between sequencing depth and locus recovery, the clean reads from each sample were mapped back to the gene sequences (exons + introns) recovered by CAPTUS using BBMAP v. 39.06 (Bushnell 2022) with settings “`kfilter=22 subfilter=15 maxindel=80`”. The average sequencing depth for each locus was then calculated using the `pileup.sh` script bundled with BBMAP.

To assess the implications of differences in locus recovery efficiency among pipelines on phylogenetic inference, we estimated coalescent species trees from the CDS recovered by each pipeline and compared their topologies and nodal supports. For CAPTUS, three sets of alignments with different paralog filtering strategies (`unfiltered`, `naive`, and `informed`) were generated using the `align` module with default settings. For HYBPIPER-BLASTX and HYBPIPER-DIAMOND, two sets of alignments were generated, one with paralogs and one without. All alignments, regardless of the pipeline, were generated using MAFFT v. 7.525 (Katoh and Standley 2013) with the automatic strategy selection mode (`--auto`) and trimmed for columns with >90% missing data using CLIPKIT v. 2.2.4 (Steenwyk et al. 2020) with the gappy mode (`-m gappy`). Coalescent species trees were estimated from each set of trimmed alignments using IQ-TREE v. 2.2.2.6 (Minh et al. 2020b) and ASTRAL-PRO v. 1.15.1.3 (Zhang et al. 2020; Zhang and Mirarab 2022) following the procedure described above, and then visualized using TOYTREE v. 2.0.5 (Eaton 2020).

## Results

### Sequencing

From our 118 library preparations for target capture, 92 (78%) worked at the first attempt. Only eight of these successful libraries needed library concentration or increased number of PCR cycles to improve their quality prior to sequencing. For the remaining 26 libraries (22%) that had to be repeated, we used material of the same herbarium voucher for 14, while we had to select a different specimen for 12. Among the repeated libraries, only two needed library concentration or increased number of PCR cycles to improve their quality prior to sequencing. Nonetheless, once the sequencing was completed, three samples had to be discarded due to insufficient data. Two of the discarded samples corresponded to replicated libraries of *Octomeles sumatrana* for which we had a replacement within the target capture batch. Only one of them, *Bambekea racemosa*, needed to be replaced by additional high-depth whole genome sequencing, which was also performed for six additional samples (for previously missing taxa). In the end, a total of 122 samples (115 target capture and 7 high-depth whole genome sequencing) yielded enough high-quality data to be used for the analyses as determined by the number of loci recovered and their completeness (Table S4).

### Read Cleaning and Assembly

The RNA-Seq data from NCBI had the largest average percentage of reads and percentage of base pairs removed with 4.7% and 9.2% respectively. The target capture data had the largest average percentage of raw reads with adaptors (14.6%) but adaptors were fully removed from all data types after cleaning (Table 1: Cleaning). Average total contig length (i.e, assembly size) after removing contigs with GC content > 60% was highest for high-depth whole genome sequencing (591.1 Mbp), followed by genome skimming (289.3 Mbp), and both target capture and RNA-Seq data with c. 44 Mbp. Among the genomic data types, the most fragmented was capture data (88.3 k contigs and N50 of 513.3 bp) while high-depth whole genome sequencing data reached the largest average N50 with 5023.7 bp. The GC content matched the expectations of the data type, being lower for genome skimming (37.5%) and high-depth whole genome sequencing (34.8%), and higher for the gene-centered datatypes target capture (40.8%) and RNA-Seq (42.6%) (Table 1: Assembly).

**TABLE 1.**
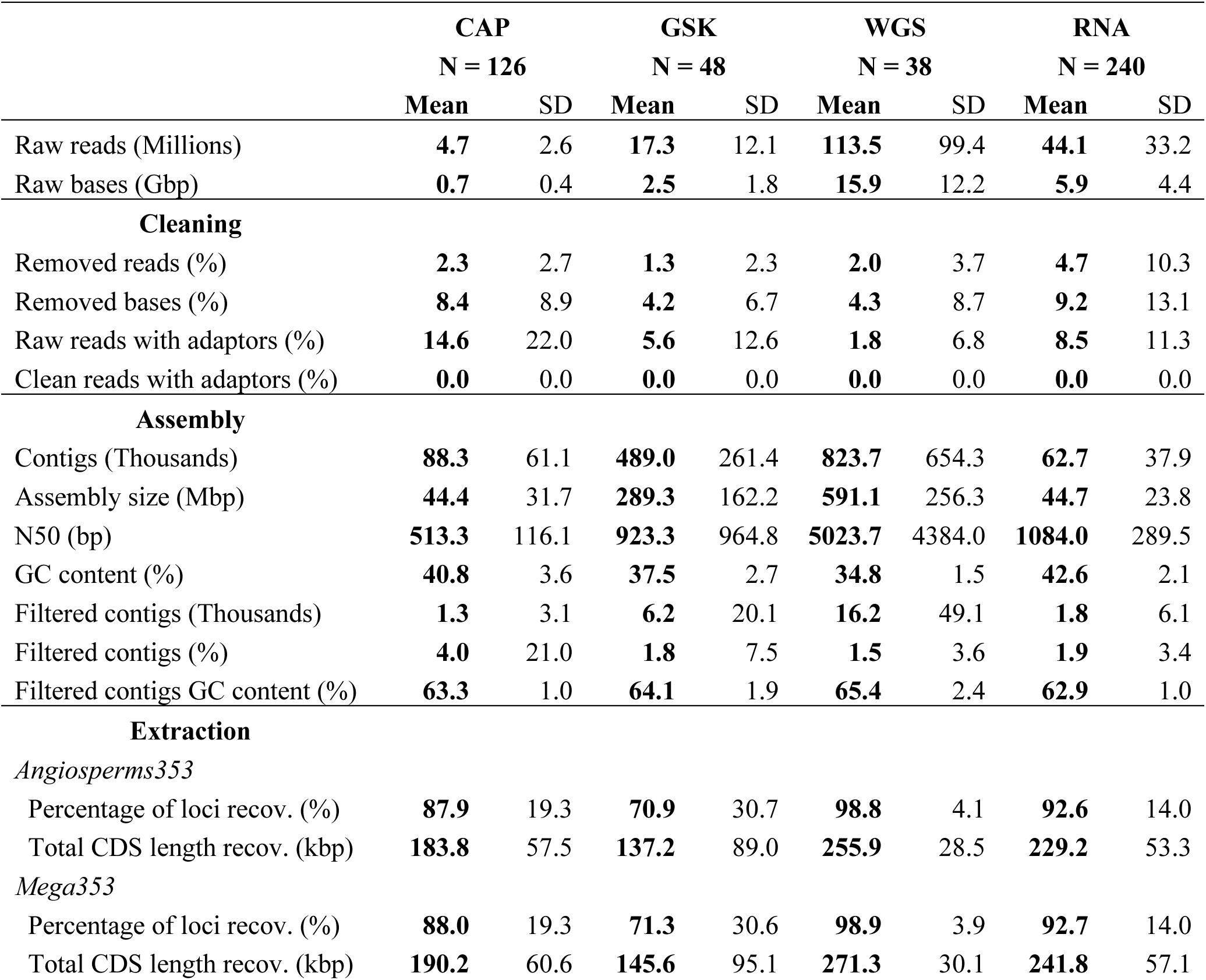
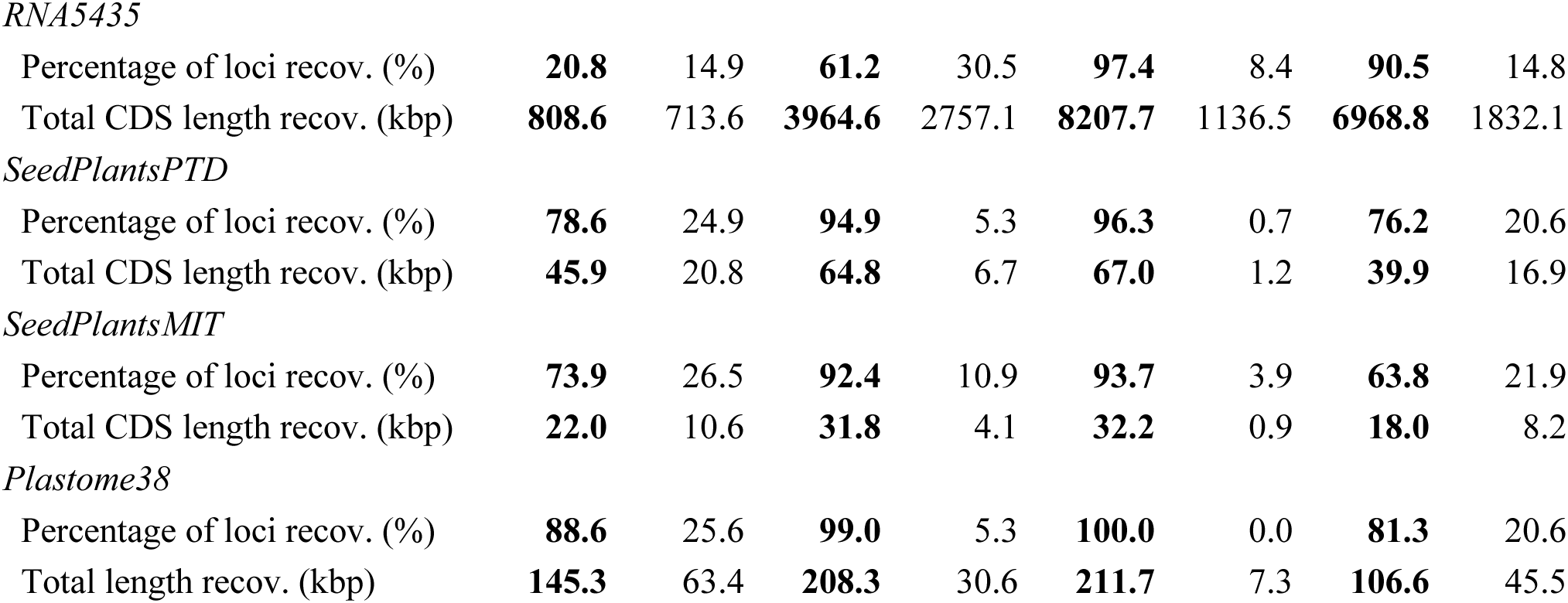
Summary statistics by data type (CAP – target capture; GSK – genome skimming; WGS – high-depth whole genome sequencing; RNA – RNA-seq).

|  | CAP<br>N = 126 |  | GSK<br>N = 48 |  | WGS<br>N = 38 |  | RNA<br>N = 240 |  |
| --- | --- | --- | --- | --- | --- | --- | --- | --- |
|  | Mean | SD | Mean | SD | Mean | SD | Mean | SD |
| Raw reads (Millions) | **4.7** | 2.6 | **17.3** | 12.1 | **113.5** | 99.4 | **44.1** | 33.2 |
| Raw bases (Gbp) | **0.7** | 0.4 | **2.5** | 1.8 | **15.9** | 12.2 | **5.9** | 4.4 |
| **Cleaning** |  |  |  |  |  |  |  |  |
| Removed reads (%) | **2.3** | 2.7 | **1.3** | 2.3 | **2.0** | 3.7 | **4.7** | 10.3 |
| Removed bases (%) | **8.4** | 8.9 | **4.2** | 6.7 | **4.3** | 8.7 | **9.2** | 13.1 |
| Raw reads with adaptors (%) | **14.6** | 22.0 | **5.6** | 12.6 | **1.8** | 6.8 | **8.5** | 11.3 |
| Clean reads with adaptors (%) | **0.0** | 0.0 | **0.0** | 0.0 | **0.0** | 0.0 | **0.0** | 0.0 |
| **Assembly** |  |  |  |  |  |  |  |  |
| Contigs (Thousands) | **88.3** | 61.1 | **489.0** | 261.4 | **823.7** | 654.3 | **62.7** | 37.9 |
| Assembly size (Mbp) | **44.4** | 31.7 | **289.3** | 162.2 | **591.1** | 256.3 | **44.7** | 23.8 |
| N50 (bp) | **513.3** | 116.1 | **923.3** | 964.8 | **5023.7** | 4384.0 | **1084.0** | 289.5 |
| GC content (%) | **40.8** | 3.6 | **37.5** | 2.7 | **34.8** | 1.5 | **42.6** | 2.1 |
| Filtered contigs (Thousands) | **1.3** | 3.1 | **6.2** | 20.1 | **16.2** | 49.1 | **1.8** | 6.1 |
| Filtered contigs (%) | **4.0** | 21.0 | **1.8** | 7.5 | **1.5** | 3.6 | **1.9** | 3.4 |
| Filtered contigs GC content (%) | **63.3** | 1.0 | **64.1** | 1.9 | **65.4** | 2.4 | **62.9** | 1.0 |
| **Extraction** |  |  |  |  |  |  |  |  |
| *Angiosperms353* |  |  |  |  |  |  |  |  |
| Percentage of loci recov. (%) | **87.9** | 19.3 | **70.9** | 30.7 | **98.8** | 4.1 | **92.6** | 14.0 |
| Total CDS length recov. (kbp) | **183.8** | 57.5 | **137.2** | 89.0 | **255.9** | 28.5 | **229.2** | 53.3 |
| *Mega353* |  |  |  |  |  |  |  |  |
| Percentage of loci recov. (%) | **88.0** | 19.3 | **71.3** | 30.6 | **98.9** | 3.9 | **92.7** | 14.0 |
| Total CDS length recov. (kbp) | **190.2** | 60.6 | **145.6** | 95.1 | **271.3** | 30.1 | **241.8** | 57.1 |
| *RNA5435* |  |  |  |  |  |  |  |  |
| Percentage of loci recov. (%) | **20.8** | 14.9 | **61.2** | 30.5 | **97.4** | 8.4 | **90.5** | 14.8 |
| Total CDS length recov. (kbp) | **808.6** | 713.6 | **3964.6** | 2757.1 | **8207.7** | 1136.5 | **6968.8** | 1832.1 |
| *SeedPlantsPTD* |  |  |  |  |  |  |  |  |
| Percentage of loci recov. (%) | **78.6** | 24.9 | **94.9** | 5.3 | **96.3** | 0.7 | **76.2** | 20.6 |
| Total CDS length recov. (kbp) | **45.9** | 20.8 | **64.8** | 6.7 | **67.0** | 1.2 | **39.9** | 16.9 |
| *SeedPlantsMIT* |  |  |  |  |  |  |  |  |
| Percentage of loci recov. (%) | **73.9** | 26.5 | **92.4** | 10.9 | **93.7** | 3.9 | **63.8** | 21.9 |
| Total CDS length recov. (kbp) | **22.0** | 10.6 | **31.8** | 4.1 | **32.2** | 0.9 | **18.0** | 8.2 |
| *Plastome38* |  |  |  |  |  |  |  |  |
| Percentage of loci recov. (%) | **88.6** | 25.6 | **99.0** | 5.3 | **100.0** | 0.0 | **81.3** | 20.6 |
| Total length recov. (kbp) | **145.3** | 63.4 | **208.3** | 30.6 | **211.7** | 7.3 | **106.6** | 45.5 |

The average GC content of the filtered contigs ranged from 59.8% to 65.4% across data types (Table 1: Assembly). The target capture data had the highest average percentage of contigs removed because of exceeding the threshold of 60% GC content (4%).

### Extraction and Alignment

Since the Angiosperms353 and the Mega353 reference targets aim to recover the same set of 353 genes, gene recovery in terms of number of loci and total CDS length was very similar for both (Table 1: Extraction). The average percentage of genes recovered across data types varied from c. 71% for genome skimming to c. 99% for high-depth whole genome sequencing data while the data type with the longest total CDS lengths is high-depth whole genome sequencing with an average of 271.3 kbp and the shortest is genome skimming with an average of 145.6 kbp. However, the Mega353 reference targets recovered at most 0.4% more loci than Angiosperms353 for genome skimming and at most 0.1% more for other data types. Similarly, the Mega353 reference targets produced total CDS lengths only c. 10 kbp longer than the Angiosperms353 reference targets on average across data types.

For the RNA5435 reference targets, target capture data had the lowest average percentage of recovered loci (20.8%, SD 14.9%) as well as the shortest average total CDS length (808.6 kbp, SD 713.6 kbp). For the rest of data types, the percentage of RNA5435 genes recovered ranges from 61.2% to 97.4% while the average total CDS length ranges from 3.96 Mbp to 8.2 Mbp. Recovery of organellar proteins was also high across data types, exceeding 73% except for mitochondrial proteins from RNA-Seq data where only 63.8% of the genes were recovered. The 38 plastome segments were successfully recovered across data types, where the minimum was 81.3% for RNA-Seq data. The total aligned ungapped length across samples was similar for Angiosperms353 (Fig. S2a) and Mega353 alignments (Fig. S2b) with c. 85% of samples with lengths between c. 200 kbp and c. 290 kbp and only c. 2.5% of samples showing less than 50 kbp aligned. As for RNA5435 (Fig. S2c) around 55% of samples had total ungapped aligned lengths between 6.0 and 8.2 Mbp, while around 20% of the samples had less than 1 Mbp aligned, corresponding mostly to target capture samples.

Based on a preliminary extraction report from CAPTUS which indicated an unusually high number of copies per DNA region in *Lemurosicyos variegata* (Cucurbitaceae), we concluded that this sample was contaminated with DNA from another plant family. Given the evolutionary distance of the contaminant, we could decontaminate the *Lemurosicyos* sequences using CAPTUS and a custom reference specific for Cucurbitaceae (Supplementary Method). We also identified the Cucurbitaceae sample *Xerosicyos perrieri* as cross-contaminated with *Ibervillea sonorae*, another Cucurbitaceae. In this case, decontamination would have implied creating a *Xerosicyos*-specific reference, which was not possible, and therefore we decided to exclude the *Xerosicyos perrieri* data from further analyses. A preliminary plastome phylogeny revealed additional samples with chloroplast contamination or lack of sufficient data for correct placement (Table S5).

### Phylogeny estimation

Coalescent species trees estimated from the set of 353 genes (using Angiosperms353 or Mega353 reference targets sets) or the RNA5435 gene trees, regardless of whether inferred from paralog-filtered or unfiltered alignments, or if estimated with FASTTREE or IQ-TREE, resulted in topologies with monophyletic and well-supported families in Cucurbitales and monophyletic and well-supported tribes in Cucurbitaceae (Fig. 2, Fig. S3, Appendix S3).

**FIGURE 2.**
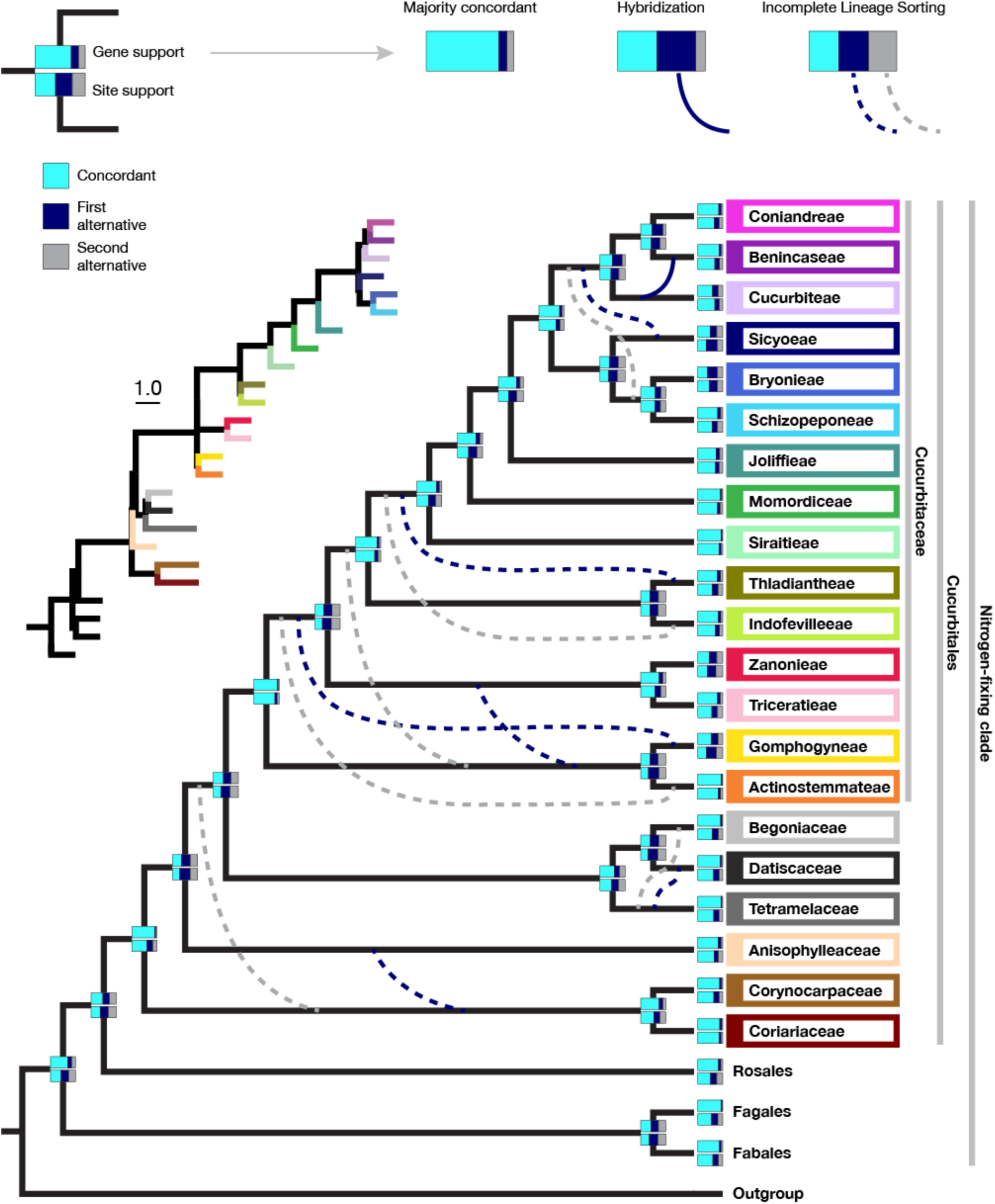
Cladogram of the Cucurbitales inferred with ASTRAL-PRO based on the RNA5435 nuclear gene trees (Fig. S5a) with families and Cucurbitaceae tribes collapsed into a single branch each. Gene support calculated with ASTRAL-PRO and site support calculated using IQ-TREE is indicated at each branch. The most frequent alternative topologies are shown with dark blue and grey dashed lines for nodes under ILS and with a continuous dark blue line for the node under hybridization. Internal branch lengths in the inset phylogram are in coalescent units.

Species trees calculated with the concatenation method for nuclear genes (Appendix S3) and for the plastome (Fig. S4) also show monophyletic families and Cucurbitaceae tribes. The holoparasitic Apodanthaceae are sister to Rafflesiaceae in Malpighiales in all inferred nuclear species trees (Fig. S3, Appendix S3), due to their holoparasitic life form, we could not recover reliable plastome data. For the mitochondrial gene regions obtained from the three Apodanthaceae samples, the results differ remarkably between genes and gene copies (Fig. 3, Table S6, Appendix S4). Focusing on what we classify “parasite DNA”, 16 mitochondrial regions support a phylogenetic position of Apodanthaceae in Malpighiales, while six regions show a placement in Cucurbitales and the remaining regions lack a well-supported phylogenetic position. For the sequences that we classify as “host DNA”, there is a strong affinity to the two host clades (28 *Apodanthes* sequences in 27 regions group with Malpighiales, 25 *Pilostyles* sequences in 20 regions group with Fabales), however, in the *Pilostyles aethiopica* data, we find five gene sequences grouping with Cucurbitales and seven of unclear affinity. This indicates that the Cucurbitales link to Apodanthaceae might be biologically meaningful and perhaps the result of an ancient host-parasite relationship that has been lost with the extinction of an ancestral lineage in the family.

**FIGURE 3.**
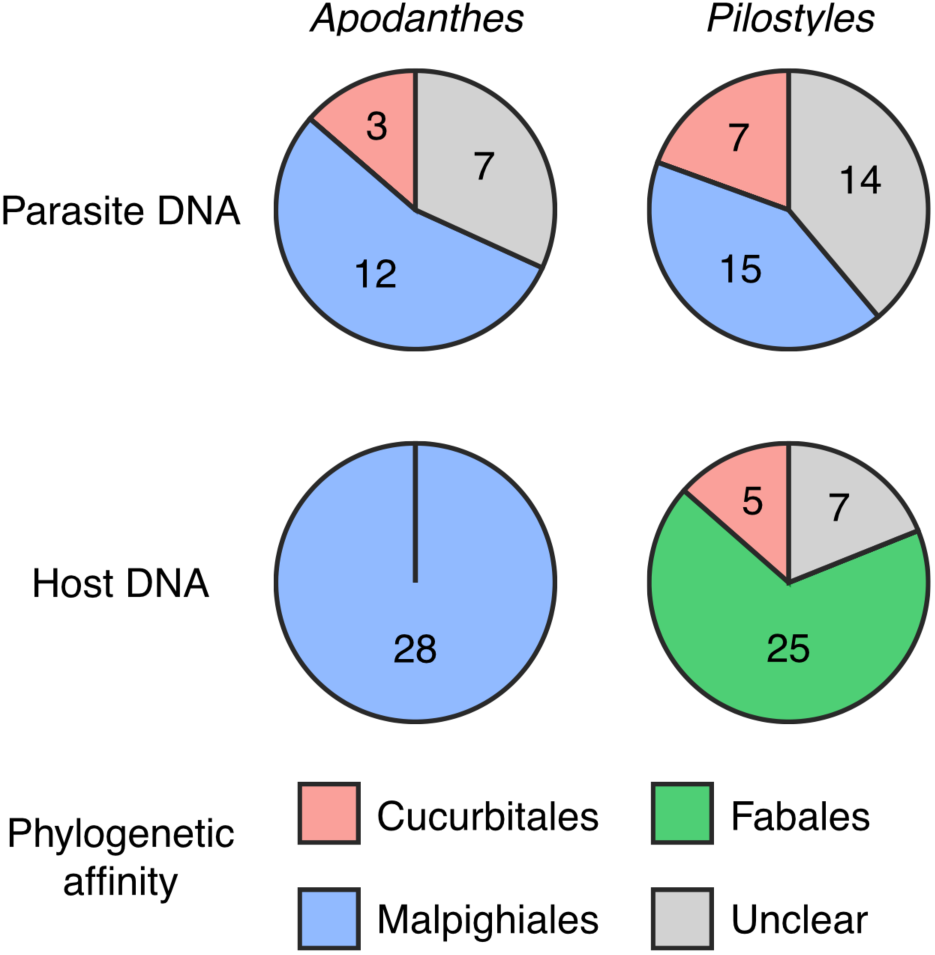
Mitochondrial affinities in the Apodanthaceae summarized by genus. The numbers represent the total number of copies found in the samples.

For the remaining Cucurbitales families, the relationships vary, particularly around the nodes with ambiguous quartet support surrounding short branches (see inset in Fig. 2).

However, the quartet frequency analyses from DISCOVISTA reveal that the apparently contradicting relationships among the major clades are all compatible with the gene tree discordance arising from ILS depicted as dashed dark blue and grey lines in Fig. 2 because the different species tree configurations observed coincide with the alternative quartet configurations suggested for the nodes in conflict (Fig. S5). The best supported alternative shows a grade, where the clade Coriariaceae + Corynocarpaceae is followed by Anisophylleaceae and by the remaining families, but there is also considerable gene tree and site support for a clade in which Anisophylleaceae is sister to a clade comprising Corynocarpaceae + Coriariaceae and the remainder of the order (Fig. 2). The woody Tetramelaceae is resolved as sister to the two herbaceous families Begoniaceae and Datiscaceae but again there is also considerable gene support for the two alternative combinations within the triplet (Fig. 2). Within Cucurbitaceae, the best supported topology is a clade comprising Actinostemmateae and Gomphogyneae as sister to the remaining tribes. Among the genera with multiple samples, only *Kedrostis* is found to be polyphyletic (Fig. S3, Appendix S3, Fig. S4). The extinct Cambodian species *Khmeriosicyos harmandii*, known only from the type collection, is confidently placed in Benincaseae as sister to *Borneosicyos* from Mount Kinabalu in the nuclear species trees (Fig. S3, Appendix S3) and in a clade with *Borneosicyos* and the southeast Asian *Solena* in the plastome tree (Fig. S4). The network analysis (Fig. S6) shows reticulation between Schizopeponeae and Sicyoeae and within and between Benincaseae and Cucurbiteae, where only the latter case is also identified as hybridization by the quartet frequency analysis (Fig. 2, Fig. S5). There is also evidence for more recent reticulation within Coniandreae and very recent within Thladiantheae (Fig. S6). The species trees resulting from the concatenated analysis of the complete plastomes (Fig. S4) also fits into the nuclear patterns of topological conflict found in Cucurbitales (Fig. 2, Fig. S6).

*Comparison with* HybPiper

The locus recovery efficiency of CAPTUS with the four different data types was evaluated against the two methods offered by HYBPIPER based on running time, number of loci recovered, and total CDS length recovered (Fig. 4a). In terms of running time, HYBPIPER-DIAMOND was the fastest, followed by CAPTUS, and then HYBPIPER-BLASTX across all tested data types, except for genome skimming, where no significant difference was found between CAPTUS and HYBPIPER-BLASTX. For the number and length of loci recovered, CAPTUS consistently recovered more loci and longer sequences than both HYBPIPER-BLASTX and HYBPIPER-DIAMOND across all tested data types. The difference in locus recovery was most pronounced in the genome skimming samples, where HYBPIPER-BLASTX and HYBPIPER-DIAMOND recovered fewer than ten loci in most samples even with the lowered coverage cutoff, while CAPTUS recovered at least 23.9 kb of CDS from 87 loci (Table S3).

**FIGURE 4.**
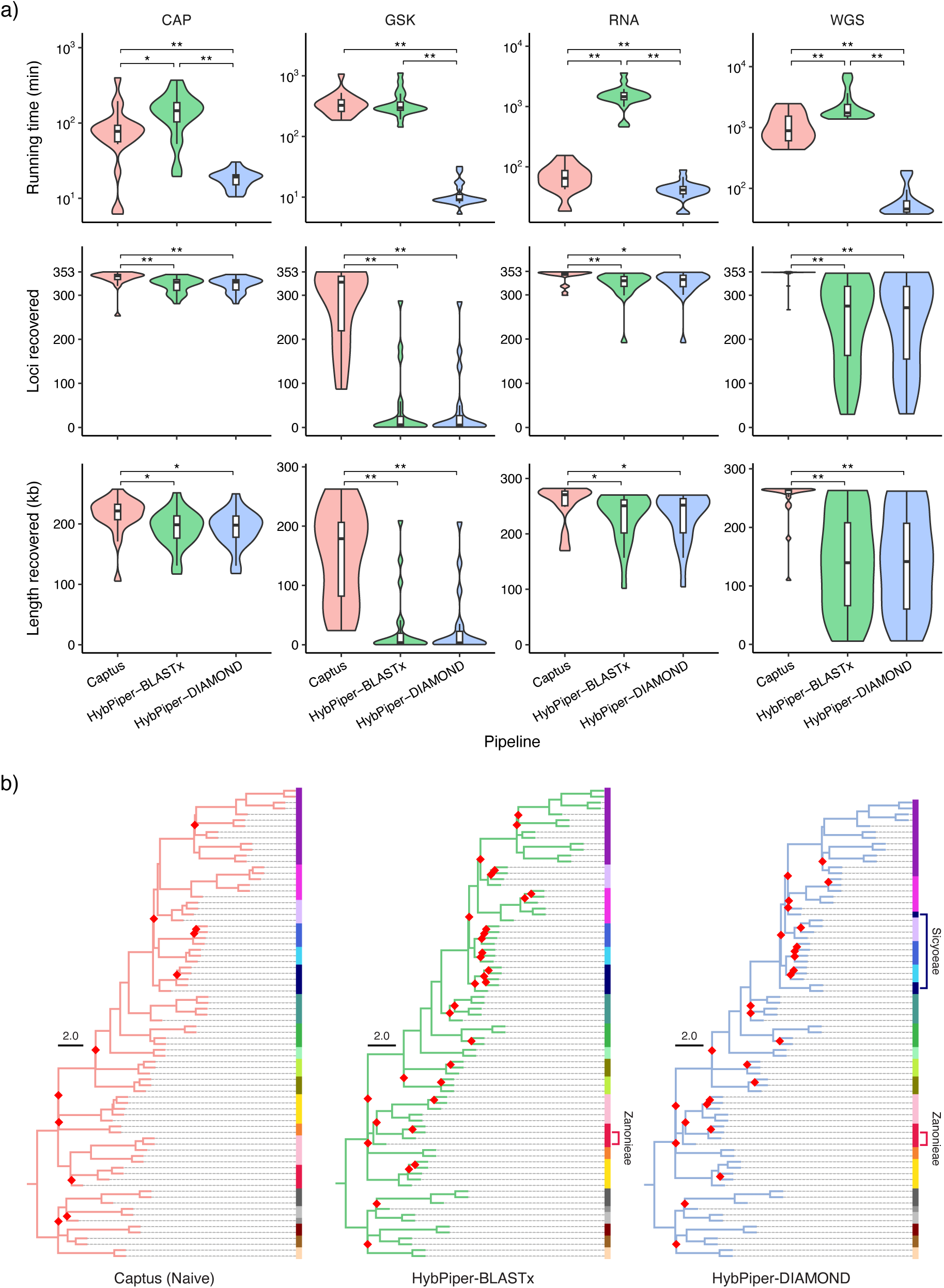
Benchmarking of CAPTUS against HYBPIPER-BLASTX and HYBPIPER-DIAMOND using a selection of 80 samples belonging to four different data types (CAP – target capture; GSK – genome skimming; RNA – RNA-seq; WGS – high-depth whole genome sequencing). **a)** Comparison of locus recovery efficiency between pipelines for each data type, evaluating the running time required for the assembly and extraction of the Angiosperms353 loci (top), the number of loci recovered (middle), and the total length of coding sequences recovered (bottom). Statistical testing was performed using the Wilcoxon rank-sum test with Bonferroni correction: * *p* < 0.05, ** *p* < 0.01. **b)** Coalescent species trees estimated from CDS alignments filtered for paralogs, resulting from each pipeline. Vertical bar along each tree color-codes the different tribes/families in the Cucurbitales. Red diamonds indicate nodes with a local posterior probability below 0.9. Tribes inferred as polyphyletic groups in the HYBPIPER trees are indicated by their names and positions. Species trees resulting from the alternative paralog filtering strategies are shown in Fig. S10. Trimmed alignments as well as the derived tree files in NEWICK format are provided in Appendix S5.

Across all tested data types, samples with lower sequencing depth tended to show greater differences in locus recovery between CAPTUS and HYBPIPER (Fig. S7). Additionally, the average sequencing depth of loci recovered only by CAPTUS was evidently lower than that of loci recovered also by HYBPIPER-BLASTX or HYBPIPER-DIAMOND (Fig. S8), highlighting the sensitivity of CAPTUS to samples with low sequencing depth. CAPTUS also showed higher sensitivity in paralog detection, recovering more paralogs than HYBPIPER-BLASTX and HYBPIPER-DIAMOND across all tested data types (Fig. S9). Collectively, these results demonstrate that although CAPTUS is slower than HYBPIPER-DIAMOND, it can extract more information for subsequent analyses with less bias across different data types, in a comparable or shorter time than HYBPIPER-BLASTX.

The differences in locus recovery efficiency between the pipelines are reflected in the results of phylogenetic inference (Fig. 4b, Fig. S10, Appendix S5). In the coalescent species trees resulting from CAPTUS, all genera, tribes, and families were recovered as monophyletic groups. In contrast, one tribe and two tribes were recovered as polyphyletic groups in the trees resulting from HYBPIPER-BLASTX and HYBPIPER-DIAMOND, respectively. These polyphylies are likely due to misplacement of *Sicyos edulis* (Sicyoeae) and *Siolmatra pentaphylla* (Zanonieae), for which HYBPIPER-BLASTX and HYBPIPER-DIAMOND recovered less than 40 loci, whereas CAPTUS recovered more than 250 loci (Table S3). In terms of nodal support, the trees resulting from CAPTUS had fewer nodes with low support (local posterior probability < 0.9) compared to those resulting from HYBPIPER-BLASTX and HYBPIPER-DIAMOND. Regardless of the pipeline, the use of paralogs or alternative filtering strategies did not significantly affect the species trees, except that alternative topologies were adopted for some of the conflicting nodes (Fig. S10).

## Discussion

Our analysis of the mixed Cucurbitales sequence data shows that CAPTUS can handle complex datasets including low-quality and contaminated sequence data and help disentangle challenging biological phenomena like ILS, hybridization, and evolution of holoparasites.

In comparison with similar existing pipelines, we find that CAPTUS is slower than HYBPIPER-DIAMOND but with our dataset recovers a higher number of more complete genes across a larger number of species than either HYBPIPER-BLASTX or HYBPIPER-DIAMOND (Fig. 4a). The obvious question arises if those additional and longer sequences recovered by CAPTUS are reliable or maybe chimeric or very low-depth sequences that could result in misleading topologies. In the case of the Cucurbitales, the tree topologies based on the CAPTUS alignments seem more reliable than the corresponding topologies from HYBPIPER. In our previous study using simulated reads at different sequencing depths, CAPTUS was shown to be capable of recovering long and accurate sequences even at low sequencing depths [Fig. 3 in (Raza et al. 2023)]. Similar to the Cucurbitales results, a higher degree of agreement among CAPTUS gene trees than for other pipelines in this earlier study [Fig. 6 in (Raza et al. 2023)] also suggests that the sequence data recovered by CAPTUS is reliable.

Turning to running time, the result of this study is inconsistent with our previous finding, in which CAPTUS was the fastest among the tested pipelines, including HYBPIPER-DIAMOND [Fig. 1 in (Raza et al. 2023)]. We hypothesize that this discrepancy can be attributed to differences in the number of target loci between the two studies: this study focused on the Angiosperms353 loci, whereas Raza et al. (2023) used a custom set of 1,180 loci. The computational cost of every step in the HYBPIPER workflow, such as distributing reads to individual loci, assembling them into contigs, and matching those contigs to reference sequences, is expected to increase proportionally to the number of target loci. In contrast, in the CAPTUS workflow, the computational cost of the `clean` and `assemble` modules remains constant regardless of the number of target loci, while only the `extract` and `align` modules are affected. This fundamental difference in workflow between the two pipelines may account for the inconsistent running time results, warranting further investigation.

The strong performance of CAPTUS in locus recovery can likely be explained by a combination of factors. First, the *de novo* assembly by MEGAHIT allows the assembly of *all* reads in the sample in contrast to HYBPIPER which only assembles subsets of prefiltered reads that match the reference loci, leading to a more restricted assembly potentially missing intronic regions and small exons and decreasing assembly contiguity in general. Second, SCIPIO’s capacity to reconstruct gene models across several contigs is essential to recover longer loci from fragmentary assemblies such as the ones resulting from target capture data and particularly genome skimming (Keller et al. 2008; Hatje et al. 2011). Finally, thanks to its improved reconstruction of short exons and intron splice sites as well as the ability to automatically reconstruct genes across multiple contigs, SCIPIO (Keller et al. 2008; Hatje et al. 2011) outperforms EXONERATE (Slater and Birney 2005), the method implemented in HYBPIPER.

The ability of CAPTUS to successfully combine different data types is confirmed by the topological results where different kinds of sequence datasets for the same species, consistently formed well-supported and stable clades (Supplement Trees). Also, by assembling the entire set of reads in a sample, CAPTUS uses off-target reads efficiently instead of excluding them a priori like HYBPIPER. Since target enrichment via DNA hybridization is an imperfect process, it commonly carries over an important amount of off-target data that can be useful for phylogenomics. Despite targeting only 353 genes, CAPTUS was able to find almost three times as many nuclear genes (c. 1130 genes on average) using our RNA5435 reference targets, as well as every plastidial and mitochondrial protein and mostly complete plastomes. Even though the number of off-target reads will greatly depend on hybridization conditions, these are usually similar for a given set of samples and a pipeline like CAPTUS can take better advantage of these other genomic regions that would remain unused otherwise.

Regarding the phylogeny estimate, our results show a well-supported nitrogen-fixing clade where Cucurbitales + Rosales are sister to Fabales + Fagales (Fig. S3, Appendix S3, Fig. S5). This topology has also been recovered by recent nuclear phylogenomic analyses (Guo et al. 2021; Kates et al. 2024; Zuntini et al. 2024) but differs from phylogenetic estimates mostly based on chloroplast data (Soltis et al. 1995; The Angiosperm Phylogeny Group 2016; Li et al. 2021) where the topology (Fabales (Rosales (Cucurbitales, Fagales) is recovered. Assuming the gain of nitrogen-fixing ability evolved in the common ancestor of this clade, the net number of independent losses inside the clade (e.g., Griesmann et al. 2018) should not be affected by this new topology, however future studies on the subject could benefit by also interpreting gains and losses according to this new topology. Within the Cucurbitales, all the currently accepted families are recovered as monophyletic groups in our analyses (Fig. 2, Fig. S3, Appendix S3, Fig. S4, Fig. S5). The relationships among families agree well with previous phylogenetic and phylogenomic studies of Cucurbitales, except for the position of the holoparasitic Apodanthaceae, where we find strong support for a position outside Cucurbitales, in contrast to the earlier results of Filipowicz and Renner (2010). We show that those earlier results can be explained by the phylogenetic signal coming from a subset of the mitochondrial genes (including the previously analyzed *matR* region, which coincidentally is the most divergent gene in the mitochondria of Apodanthaceae), while *most* mitochondrial genes group with Malpighiales. This affinity of Apodanthaceae with Malpighiales is further strongly supported by the nuclear data and in agreement with our recent result based on a 58% sampling of the 13,600 genera of angiosperms (Zuntini et al. 2024). However, the results of the CAPTUS analyses indicate a highly supported sister group relationship between the Apodanthaceae and the Rafflesiaceae (Fig. S3, Appendix S3, Fig. S5) while in our previous angiosperm-wide analysis (Zuntini et al. 2024) this is not clear. The results of Alzate et al. (2024) with a reduced taxonomic sampling seem to support the inclusion of Apodanthaceae in Rafflesiaceae but a deeper phylogenomic analysis of Malpighiales that takes advantage of the increased taxon sampling in Zuntini et al. (2024) and our increased gene sampling might be required for a robust test of these relationships.

Although there is strong evidence of horizontal gene transfer between parasites and hosts in angiosperms, where the parasites can act either as donors or recipients mainly via DNA-mediated transfer events, such transfers are considered “recent” and usually functional and pseudogenized copies persist in the recipient (Mower et al. 2010). The pattern observed in this study for the subset of mitochondrial genes with a close relationship between Apodanthaceae and Cucurbitales does not show any of these characteristics and therefore poses questions worth investigating with a deeper analysis.

For the autotrophic families we find most support for a grade where the clade Coriariaceae + Corynocarpaceae is followed by Anisophylleaceae, a clade with the triplet Tetramelaceae plus Begoniaceae and Datiscaceae, and by Cucurbitaceae (Fig. 2, Fig. S3a, Fig. S5). Earlier studies placed Anisophylleaceae as sister to all other except Apodanthaceae (Zhang et al. 2006; Schaefer and Renner 2011b; Zuntini et al. 2024). Even though our analyses show the highest support for the clade Coriariaceae + Corynocarpaceae as sister to the remaining Cucurbitales, there is almost equal support among gene trees for Anisophylleaceae as sister to the rest (as in previous studies) as well as for a clade (Anisophylleaceae (Coriariaceae, Corynocarpaceae)) as sister to the rest of families in the order (Fig. 2, Fig. S3a, Fig. S5). For Datiscaceae, Zhang et al. (2006) and Schaefer and Renner (2011b) also found a sister group relationship to Begoniaceae, albeit with low support. Here, we find that such a clade has the highest gene tree support but the alternatives Datiscaceae sister to Tetramelaceae [recovered by Zuntini et al. (2024)] and Datiscaceae sister to Begoniaceae + Tetramelaceae also receive almost equal support (Fig. 2, Fig. S5). In the network (Fig. S6), the two dioecious families Datiscaceae + Tetramelaceae form a clade, which is sister to the monoecious Begoniaceae.

Tribal relationships within Cucurbitaceae match well the phylotranscriptomics results of Guo et al. (2020). The four conflicting nodes identified in Bellot et al. (2020) concerning the position of *Luffa*, *Hodgsonia*, *Bryonia*, and *Indofevillea* are stable when coalescent and concatenated phylogeny estimates are compared (Fig. S3, Appendix S3): *Indofevillea* is placed as sister to the Southeast Asian Thladiantheae; the sponge gourds (*Luffa*) are sister to all other Sicyoeae; the Asian *Hodgsonia* with the Neotropical Sicyoeae are sister to *Trichosanthes*; and Bryonieae plus Schizopeponeae are sister to Sicyoeae. Looking at the gene tree analyses, however, the considerable number of trees supporting alternative positions of those lineages is evident (Fig. 2, indicated in dark blue and grey dashed lines).

CAPTUS can reduce the amount of phylogenetic conflict introduced by the assembly and loci retrieval methods as it tends to show a higher degree of agreement between gene trees and the species tree than other methods (Raza et al. 2023). This provides the opportunity for careful examination of topological conflict and its causes, allowing us to show the common occurrence of ILS during the evolution of Cucurbitales as well as the detection of a deep reticulation event, which seem to be prevalent across the angiosperms (Stull et al. 2023). While the DiscoVista visualization can identify a putative hybridization event if it happened between the branches of a quartet, the detection of hybridization events between non-adjacent branches in the species tree will require more formal tests that address reticulation directly (Solís-Lemus et al. 2017; Wen et al. 2018; Kong et al. 2025; Kolbow et al.). The comparison of the gene tree frequencies obtained from the 353 captured regions (Angiosperms353, or the taxonomically expanded Mega353) with the frequencies obtained from the RNA5435 regions shows stable 1/3:1/3:1/3 ratio for nodes showing incomplete lineage sorting (e.g., nodes 14, 24, 29, 32, and 41 in Fig. S5a) or 1/2:1/2:0 ratio for nodes of hybrid origin (e.g., node 7 in Fig. S5a). This indicates that adding more genomic regions is unlikely to change the family and tribal level relationships in Cucurbitales found in our study. Future work should focus on the more recent evolutionary history of the clade and phylogenetic patterns within Cucurbitaceae genera.

In conclusion, we show that our new pipeline can handle a complex phylogenomic analysis in a very efficient way, maximizing the amount of recovered data in a reasonable time frame while maintaining outputs organized across a large number of samples and loci. The clustering capability of CAPTUS enables its application not only to seed plants but to any taxonomic group, even those where a reference set of orthologous loci has not yet been developed. Thus, CAPTUS can be used as a user-friendly universal tool for the assembly of phylogenomic datasets, even with mixed data of different origins and degraded or contaminated samples.

Data Availability

All the original sequence data is available in Genbank (see Table S1 for accession numbers), Captus commands to reproduce the results are provided in the Material and Methods section. Supplements provided as a .zip file through bioRxiv.

## Acknowledgements

We are grateful to the curators of the herbarium of Royal Botanic Gardens, Kew (K), the herbarium of Muséum National d’Histoire Naturelle in Paris (P), and the herbarium of the Botanische Staatssammlung München (M) for permission to extract DNA from selected specimens. We also thank A. Tellier (TUM) for access to the Population Genetics HPC and P.S. Soltis, D.E. Soltis, B. Faircloth and two anonymous reviewers for comments that help to improve the manuscript. The study was funded by the German Science Foundation DFG - SCHA 1875/4-2 within SPP 1991 Taxon-Omics (to HS), and by grants from the Calleva Foundation to the Plant and Fungal Trees of Life Project (PAFTOL) at the Royal Botanic Gardens, Kew.

## Supplementary Material

SUPPLEMENTARY METHOD. Decontamination process for target capture sample *Lemurosicyos variegata*.

TABLE S1. List of samples sequenced for this study.

TABLE S2. List of samples with public data used in this study.

TABLE S3. Per-sample comparison of running time, number of loci recovered, total CDS length recovered, and average number of paralogs per recovered locus among CAPTUS, HYBPIPER-BLASTX, and HYBPIPER-DIAMOND.

TABLE S4. Summary statistics per sample at each analysis step.

TABLE S5. Samples lacking sufficient chloroplast data for phylogenetic estimation or with chloroplast contamination.

TABLE S6. Mitochondrial affinities in the Apodanthaceae. The number of copies found is indicated next to the letter abbreviations for the species. A=*Apodanthes caseariae*, Pa=*Pilostyles aethiopica*, Pt=*Pilostyles thurberi*.

FIGURE S1. The CAPTUS output formats, **a)** outputs available when a protein extraction (`NUC, PTD, MIT`) is performed, **b)** outputs available when a miscellaneous DNA extraction (`DNA, CLR`) is performed.

FIGURE S2. Total aligned ungapped length per sample using different nuclear reference targets sets, **a)** Angiosperms353, **b)** Mega353, **c)** RNA5435.

FIGURE S3. Coalescent species trees estimated with ASTRAL-PRO using gene trees estimated by IQ-TREE on alignments derived from different reference targets sets (RNA5435, Angiosperms353, Mega353) and paralog filtering strategies (`unfiltered`, `informed`, `naive`).

FIGURE S4. Species tree estimated by IQ-TREE on the concatenated alignments derived from the Plastome38 reference targets set.

FIGURE S5. DISCOVISTA analyses of relative quartet frequencies imposed on the coalescent ASTRAL-PRO topology shown in Fig. S3a using sets of gene trees derived from the different reference target sets (RNA5435, Angiosperms353, Mega353) calculated by different programs (IQ-TREE, FASTTREE) and filtered for paralogs (`informed`, `naive`).

FIGURE S6. Phylogenetic network estimate for the Cucurbitales calculated with SPLITSTREE using the Neighbor Net algorithm on the concatenated alignment of the Angiosperms353 alignments which were filtered using the informed filter of CAPTUS.

FIGURE S7. Relationship between sequencing depth and locus recovery for each datatype. FIGURE S8. Distribution of sequencing depth of loci recovered by CAPTUS. Loci are classified according to whether they were recovered only by CAPTUS, or also by HYBPIPER-BLASTX or HYBPIPER-DIAMOND. Vertical dashed lines indicate the median value for each series.

FIGURE S9. Comparison of paralog sensitivity between pipelines for each data type. Statistical testing was performed using the Wilcoxon rank-sum test with Bonferroni correction: * *p* < 0.05, ** *p* < 0.01.

FIGURE S10. Coalescent species trees of 80 samples selected for the benchmarking, estimated from CDS alignments filtered or unfiltered for paralogs. Vertical bar along each tree color-codes the different tribes/families in the Cucurbitales. Red diamonds indicate nodes with a local posterior probability below 0.9. Tribes inferred as polyphyletic groups in the HYBPIPER trees are indicated by their names and positions.

APPENDIX S1. Plastome segments used as reference targets provided as a file in FASTA format `Plastome38.fasta`

APPENDIX S2. Newly found nuclear putative homologs used as reference targets provided as a file in FASTA format `RNA5435.fasta`

APPENDIX S3. Estimated species trees in NEWICK format.

APPENDIX S4. Alignments and trees of the mitochondrial regions in NEXUS format.

APPENDIX S5. Benchmarking data: Trimmed gene alignments in FASTA format and their estimated gene and species trees in NEWICK format.

APPENDIX S6. Reference targets sets used in the decontamination of *Lemurosicyos variegata* (Supplementary Method).

