## Supplementary material for "A novel phylogenomics pipeline reveals extensive topological conflict in the evolution of the angiosperm order Cucurbitales": FigS3.pdf

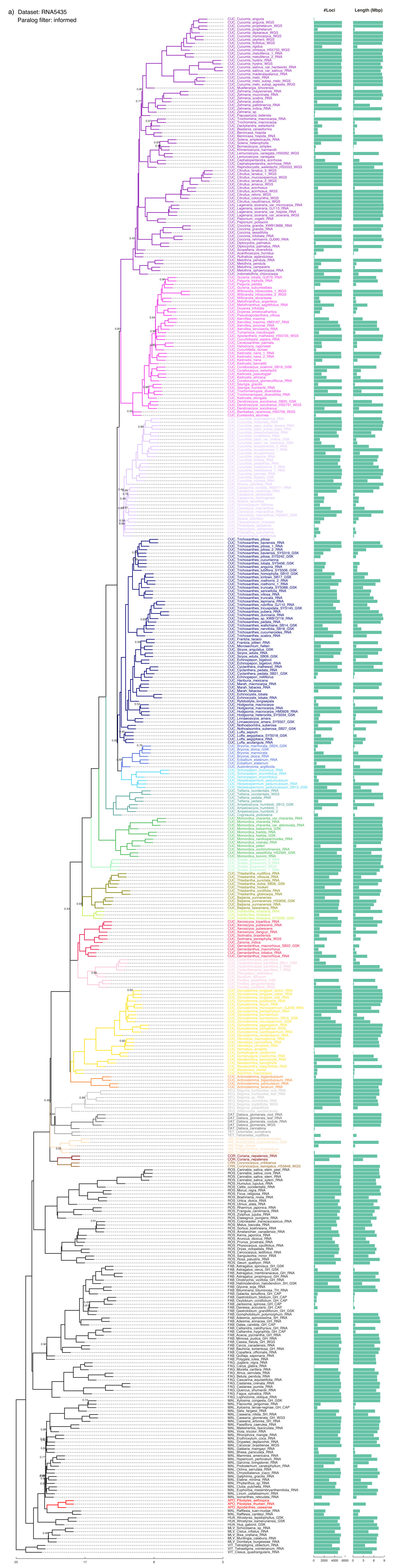

a) Dataset: RNA5435
Paralog filter: informed
#Loci
Length (Mbp)
25
17
8
0
0
2000
4000
6000
0
3
6
9

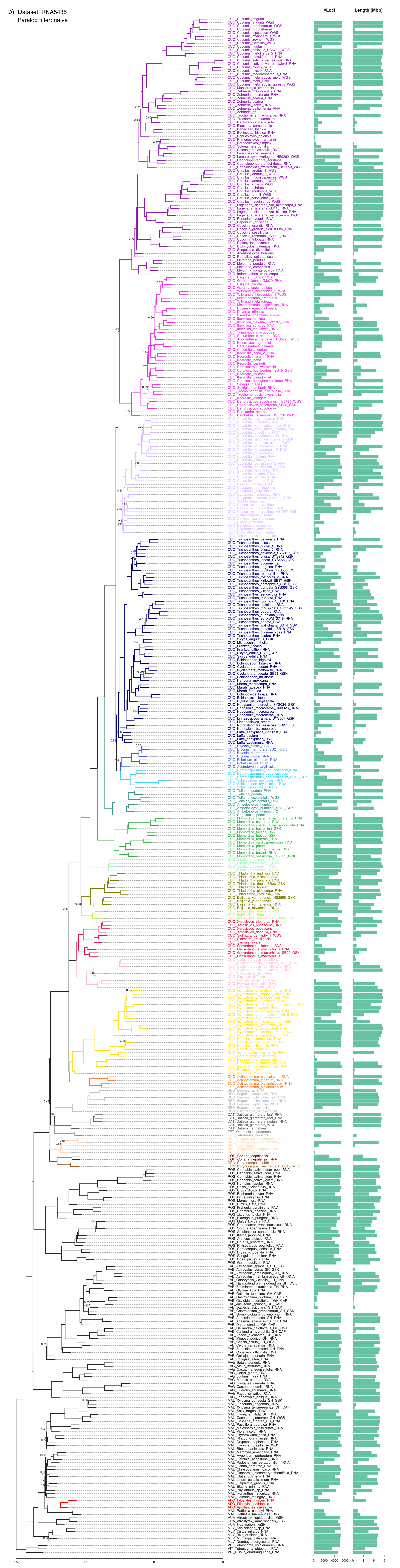

b) Dataset: RNA5435
Paralog filter: naive
#Loci
Length (Mbp)
25
17
8
0
0 2000 4000 6000
0 3 6 9

c) Dataset: RNA5435
Paralog filter: unfiltered

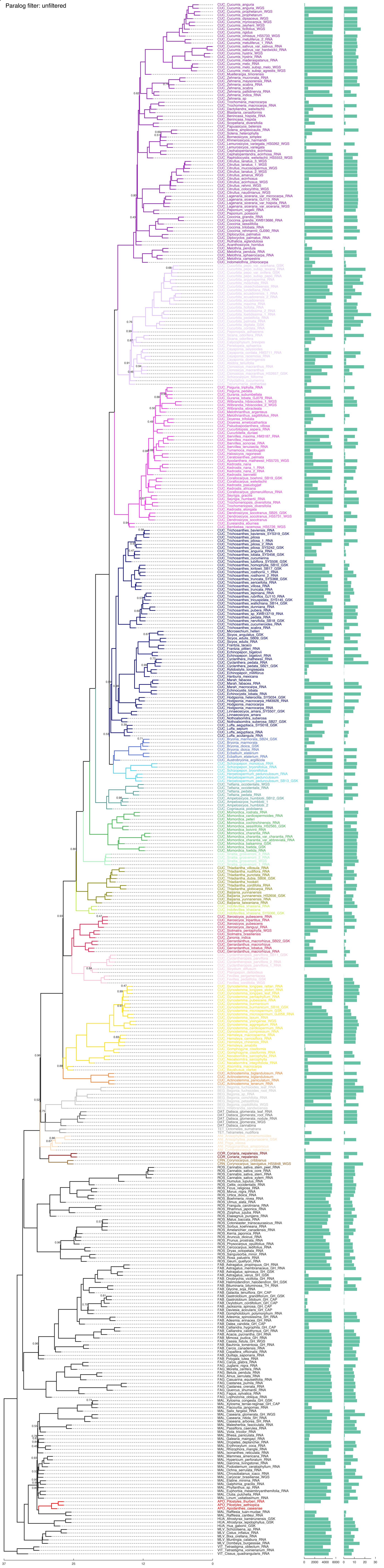

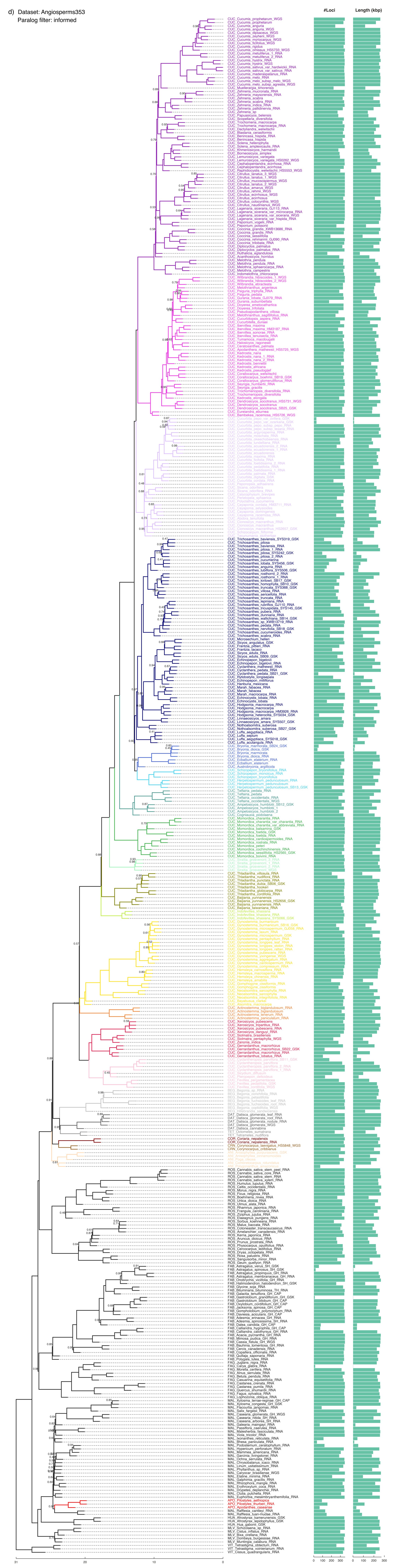
d) Dataset: Angiosperms353
Paralog filter: informed
#Loci
Length (kbp)

e) Dataset: Angiosperms353
Paralog filter: naive

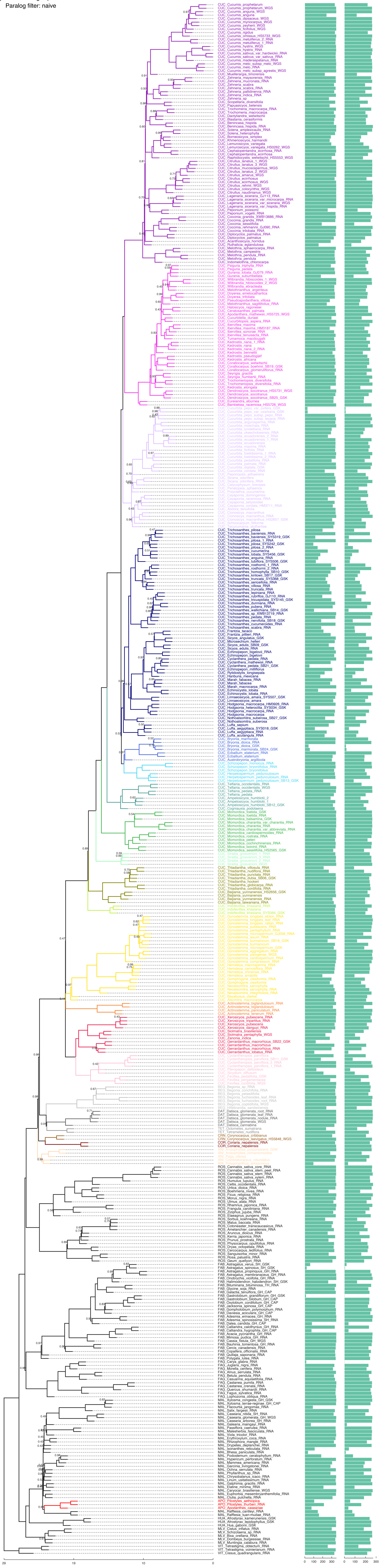

f) Dataset: Angiosperms353
Paralog filter: unfiltered

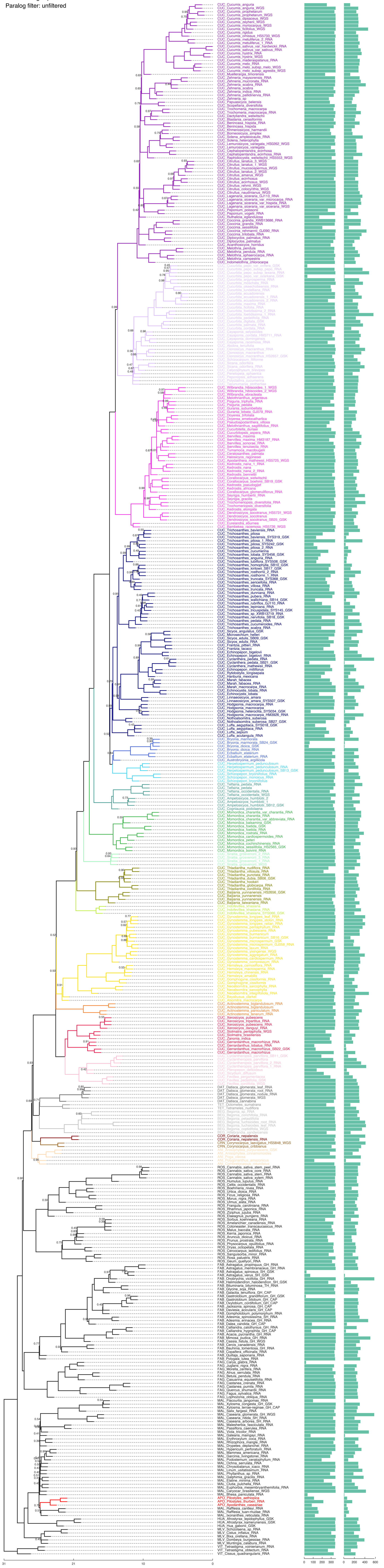

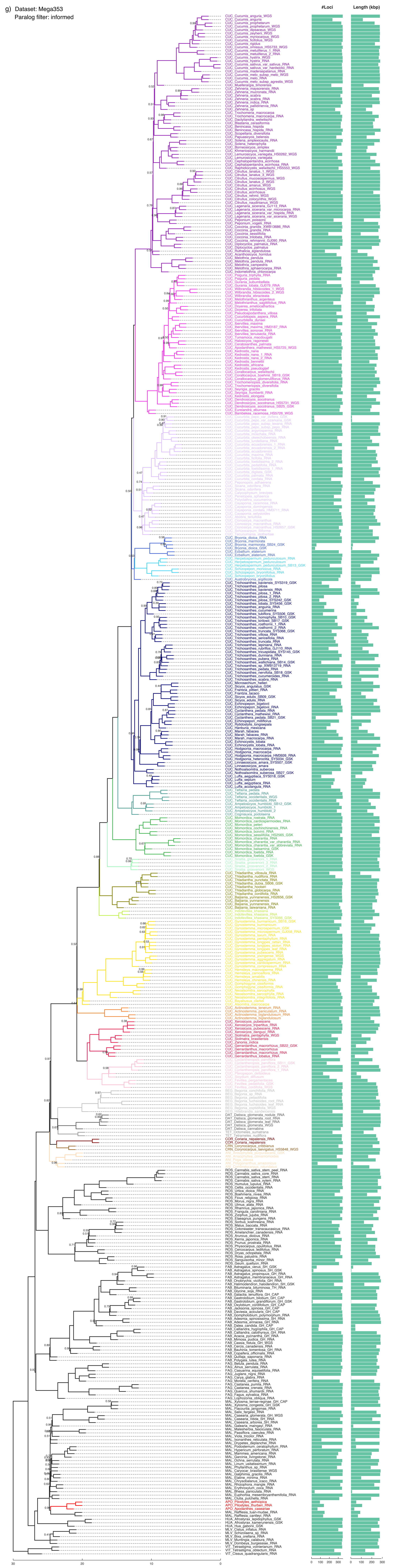
g) Dataset: Mega353
Paralog filter: informed
#Loci
Length (kbp)

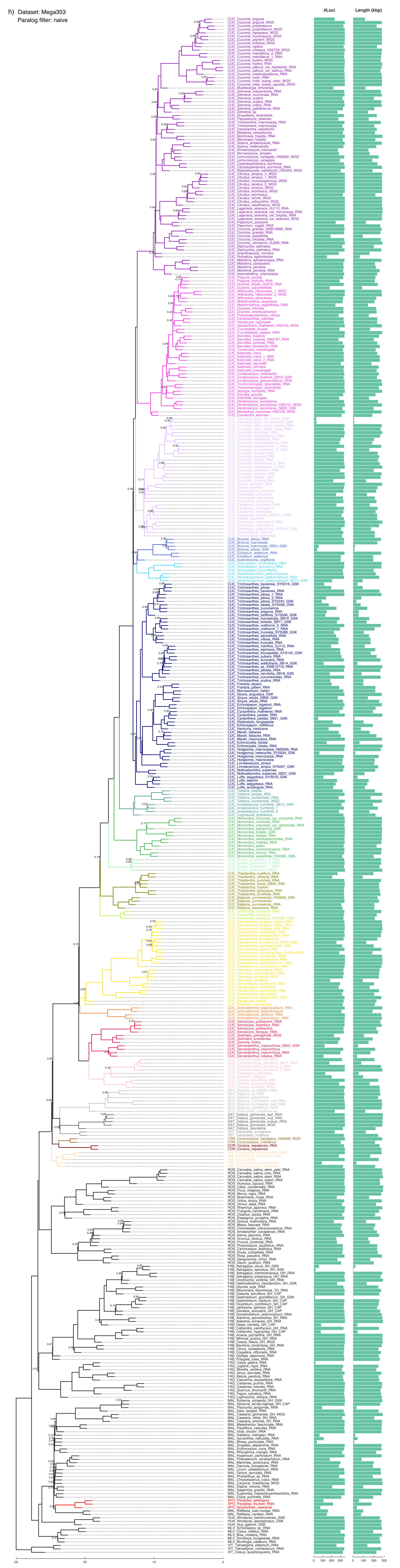
h) Dataset: Mega353
Paralog filter: naive
#Loci
Length (kbp)

i) Dataset: Mega353
Paralog filter: unfiltered

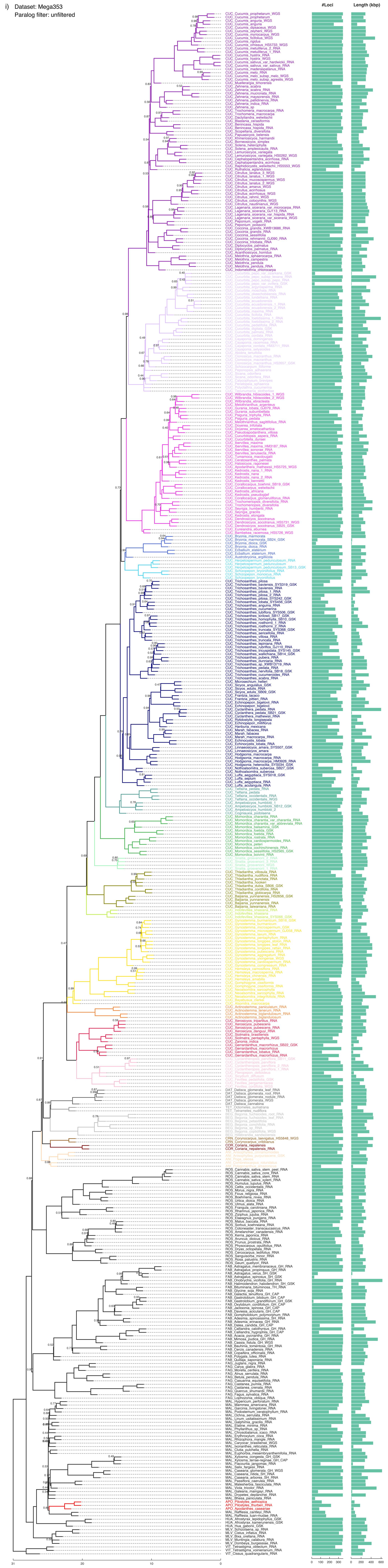
