## Supplementary material for "A novel phylogenomics pipeline reveals extensive topological conflict in the evolution of the angiosperm order Cucurbitales": FigS5.pdf

a) Method: IQ-TREE

Dataset: RNA5435

Paralog filter: informed

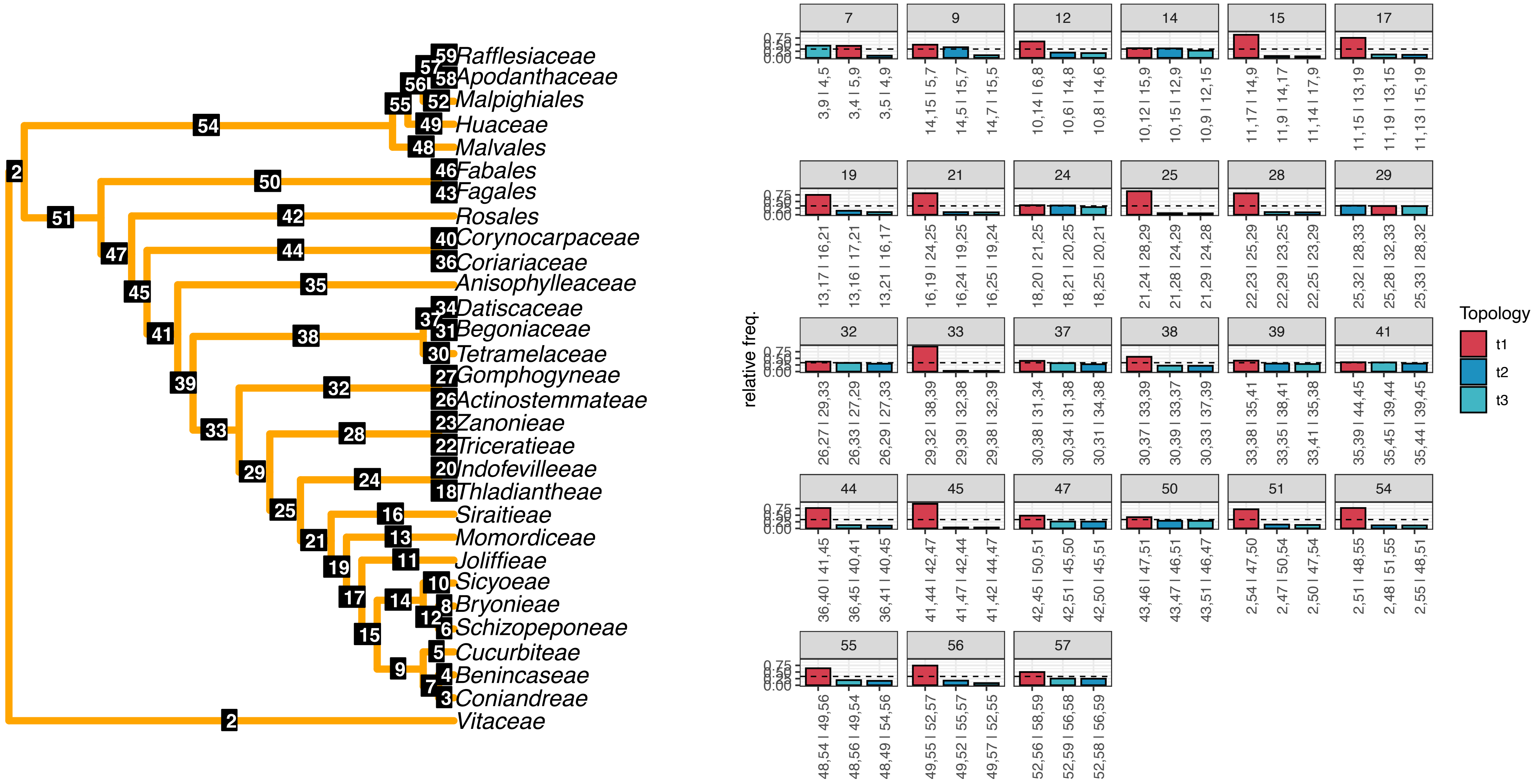

b) Method: IQ-TREE
Dataset: RNA5435
Paralog filter: naive

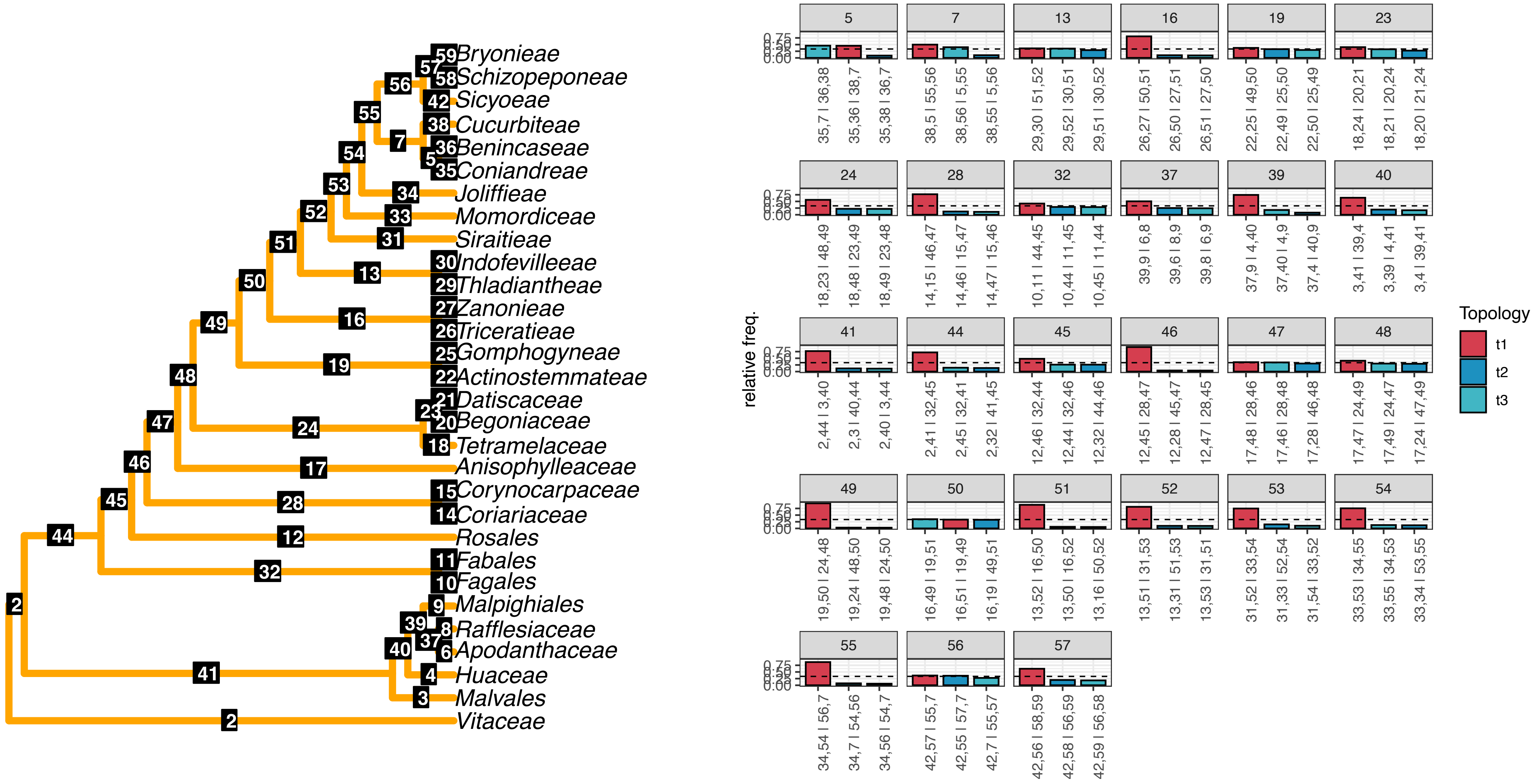

c) Method: IQ-TREE
Dataset: Angiosperm353
Paralog filter: informed

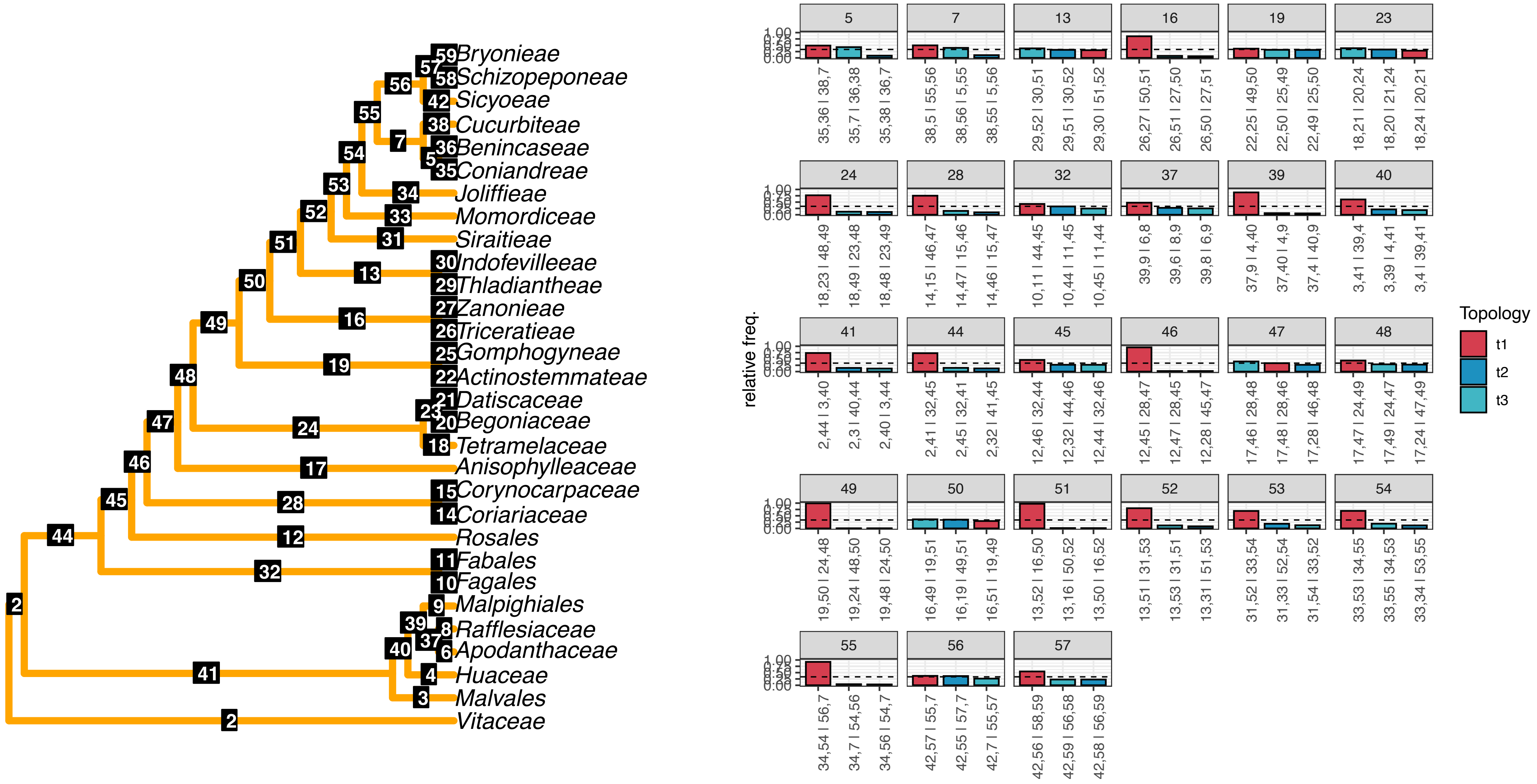

d) Method: IQ-TREE

Dataset: Angiosperm353

Paralog filter: naive

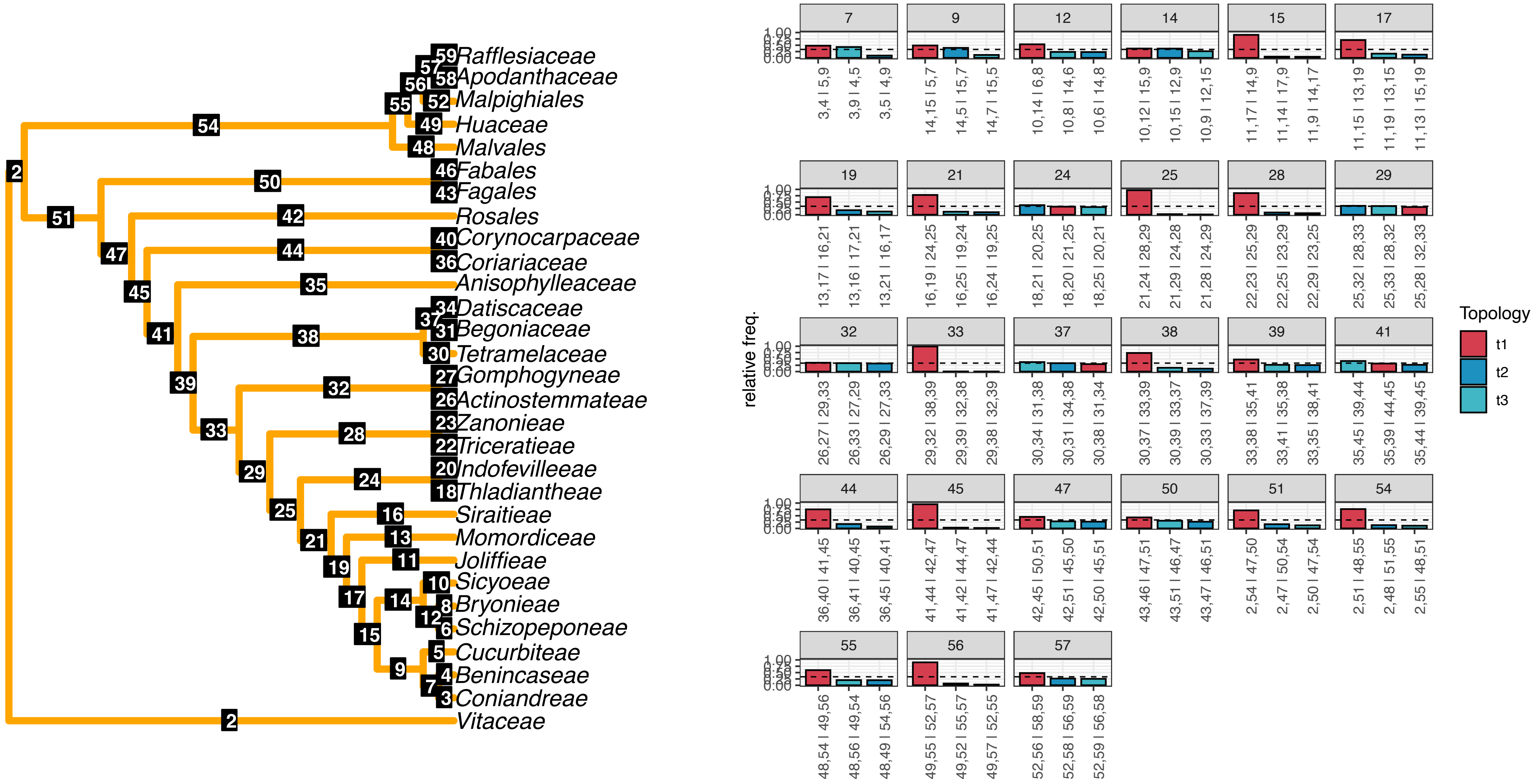

e) Method: IQ-TREE

Dataset: Mega353

Paralog filter: informed

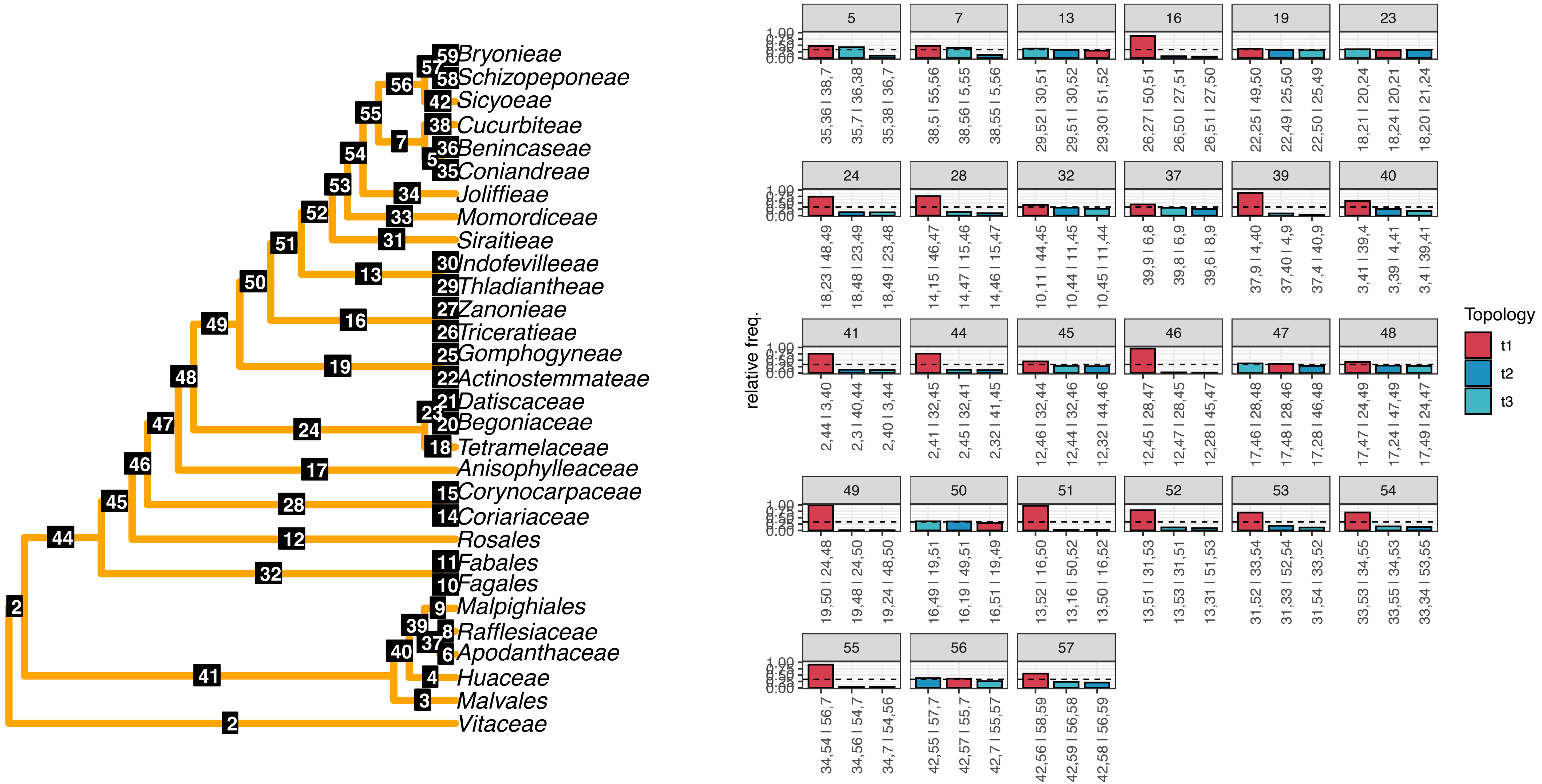

f) Method: IQ-TREE
Dataset: Mega353
Paralog filter: naive

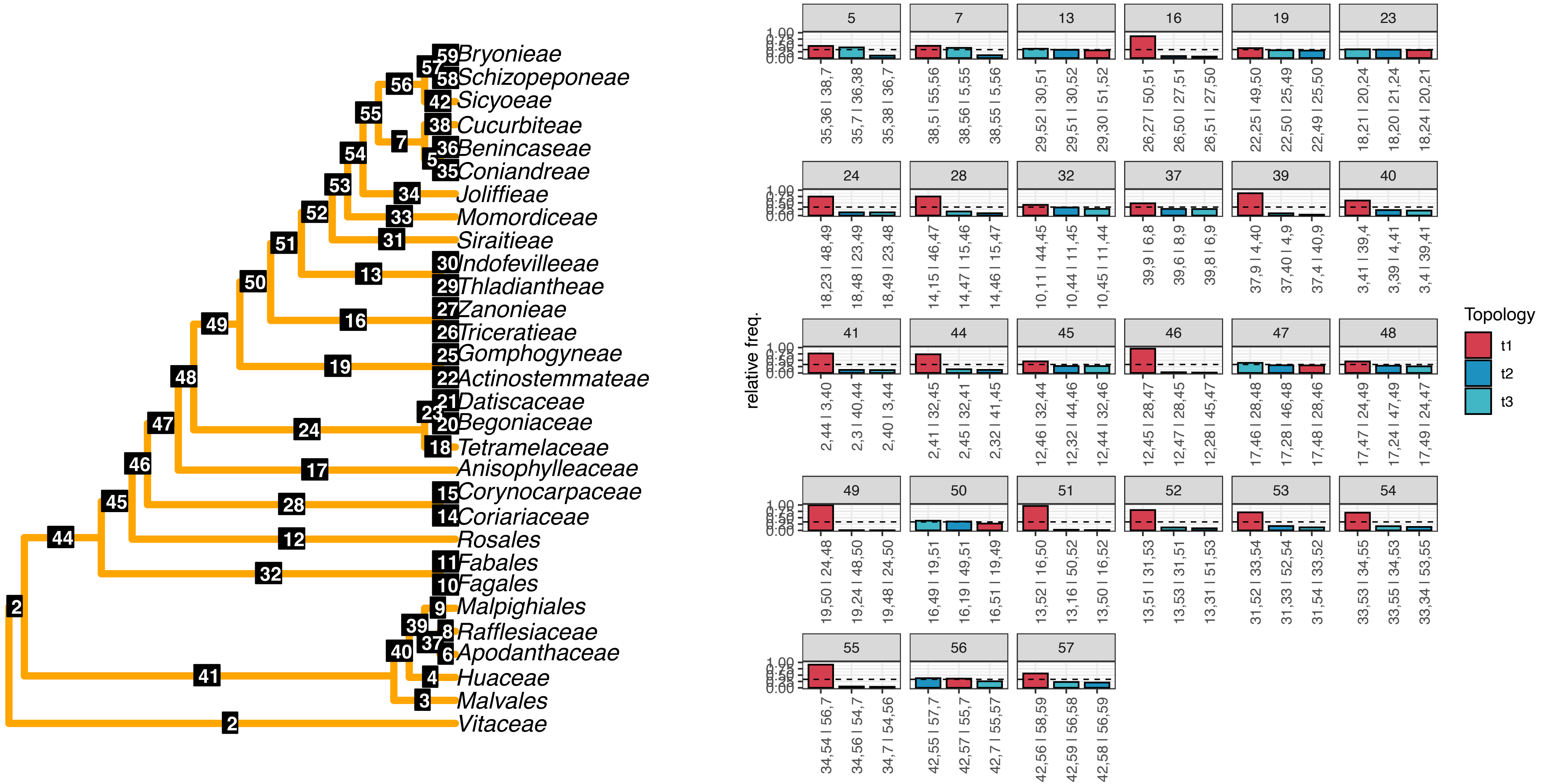

g) Method: FastTree
Dataset: RNA5435
Paralog filter: informed

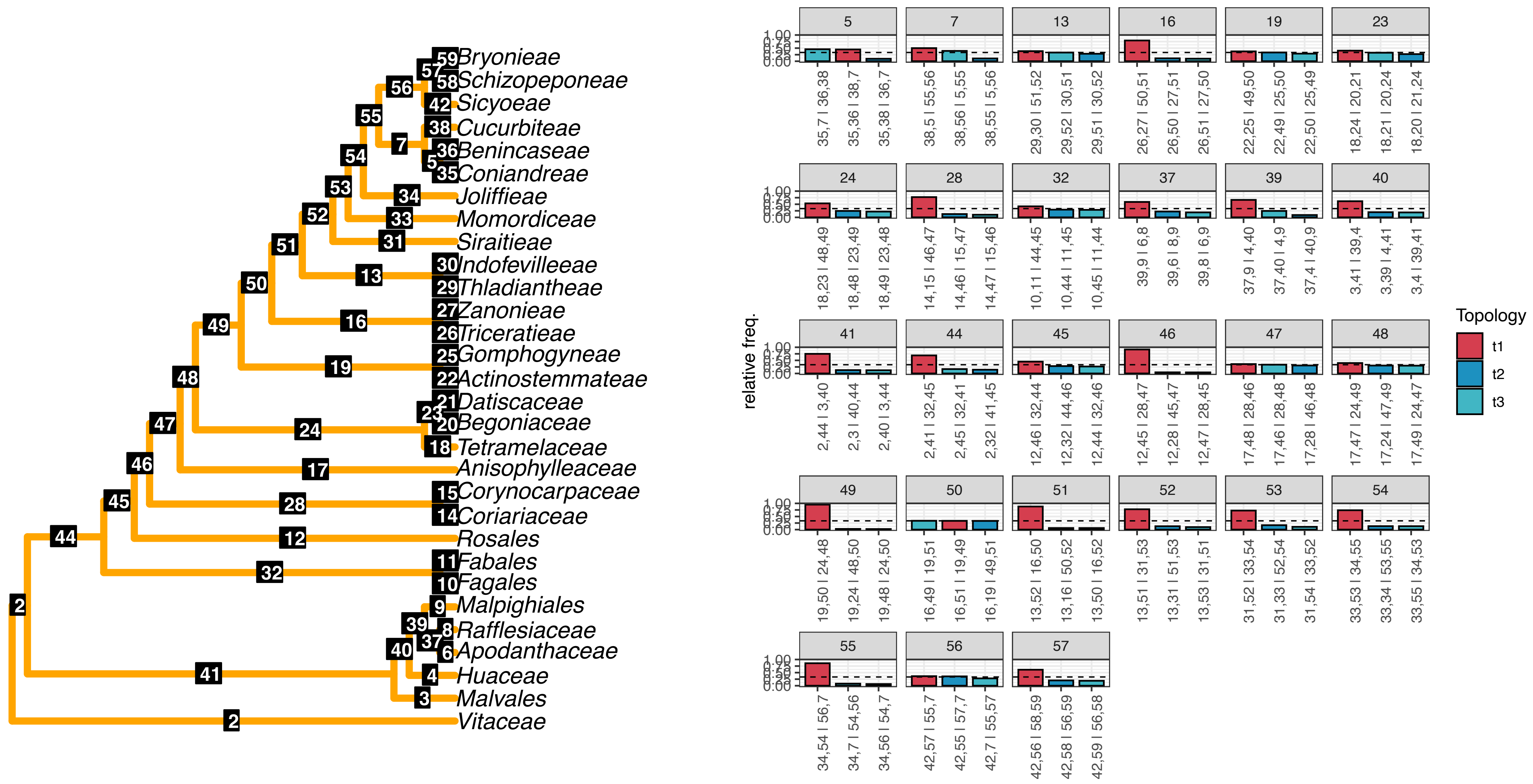

h) Method: FastTree

Dataset: RNA5435

Paralog filter: naive

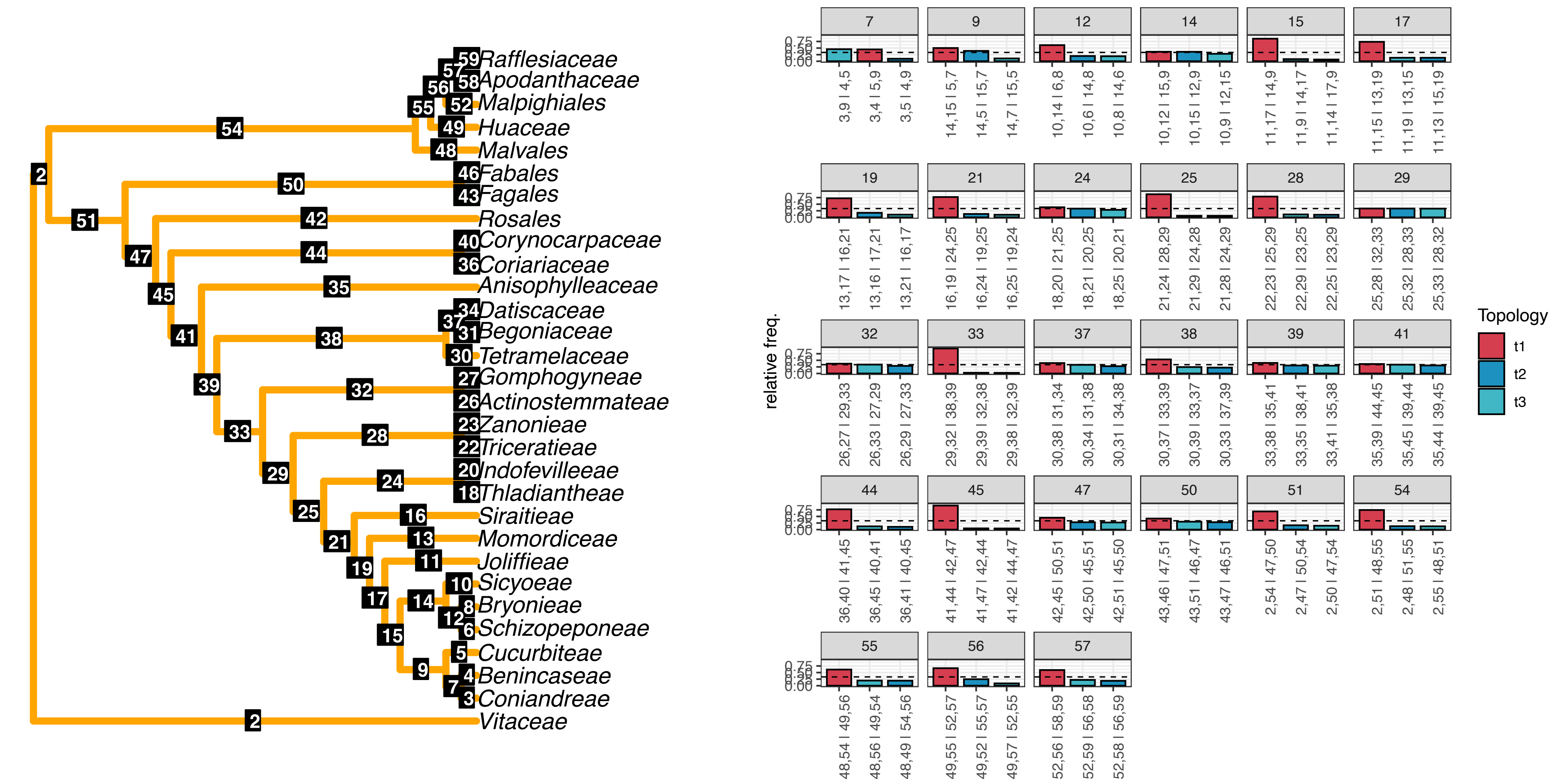

i) Method: FastTree
Dataset: Angiosperm353
Paralog filter: informed

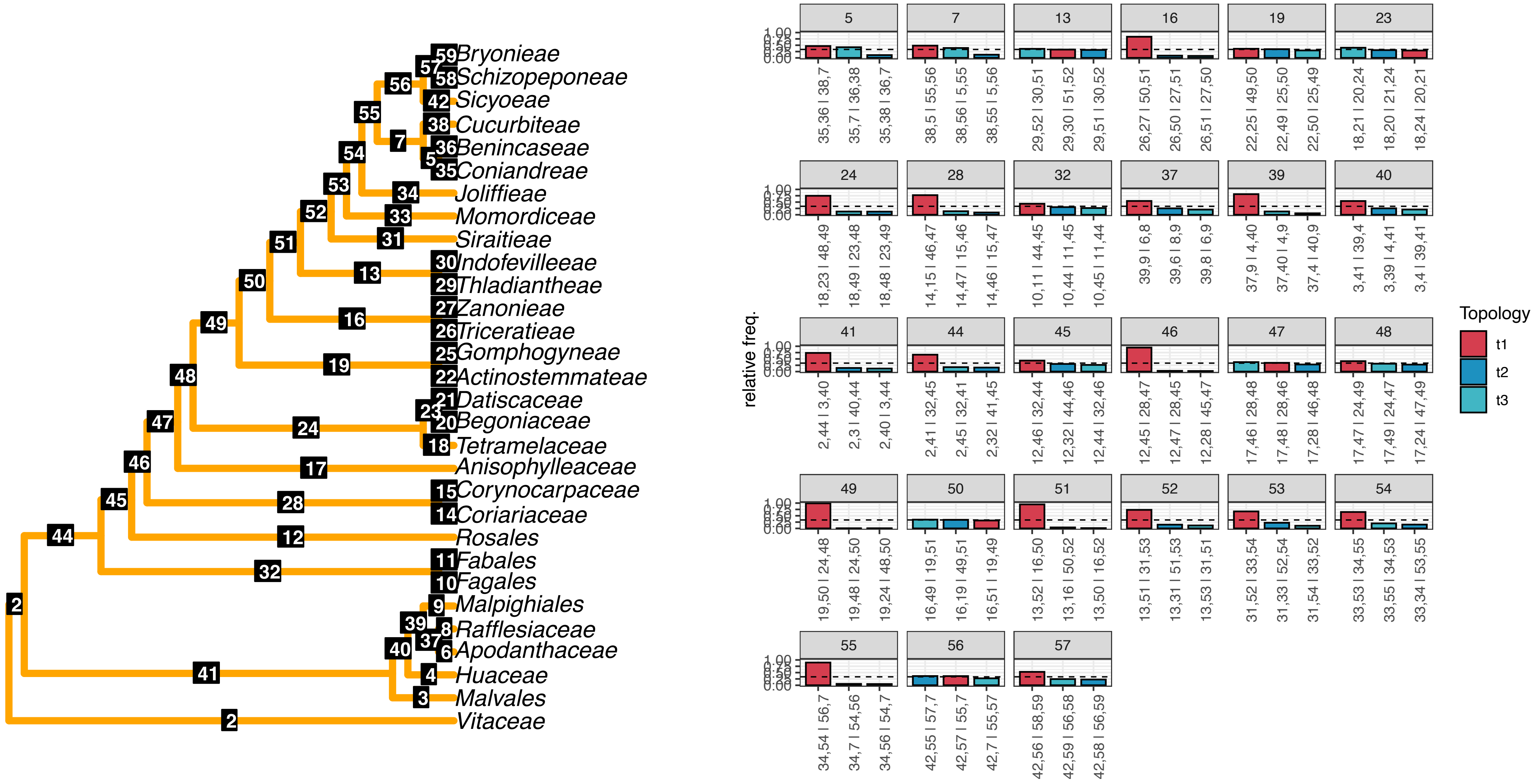

j) Method: FastTree
Dataset: Angiosperm353
Paralog filter: naive

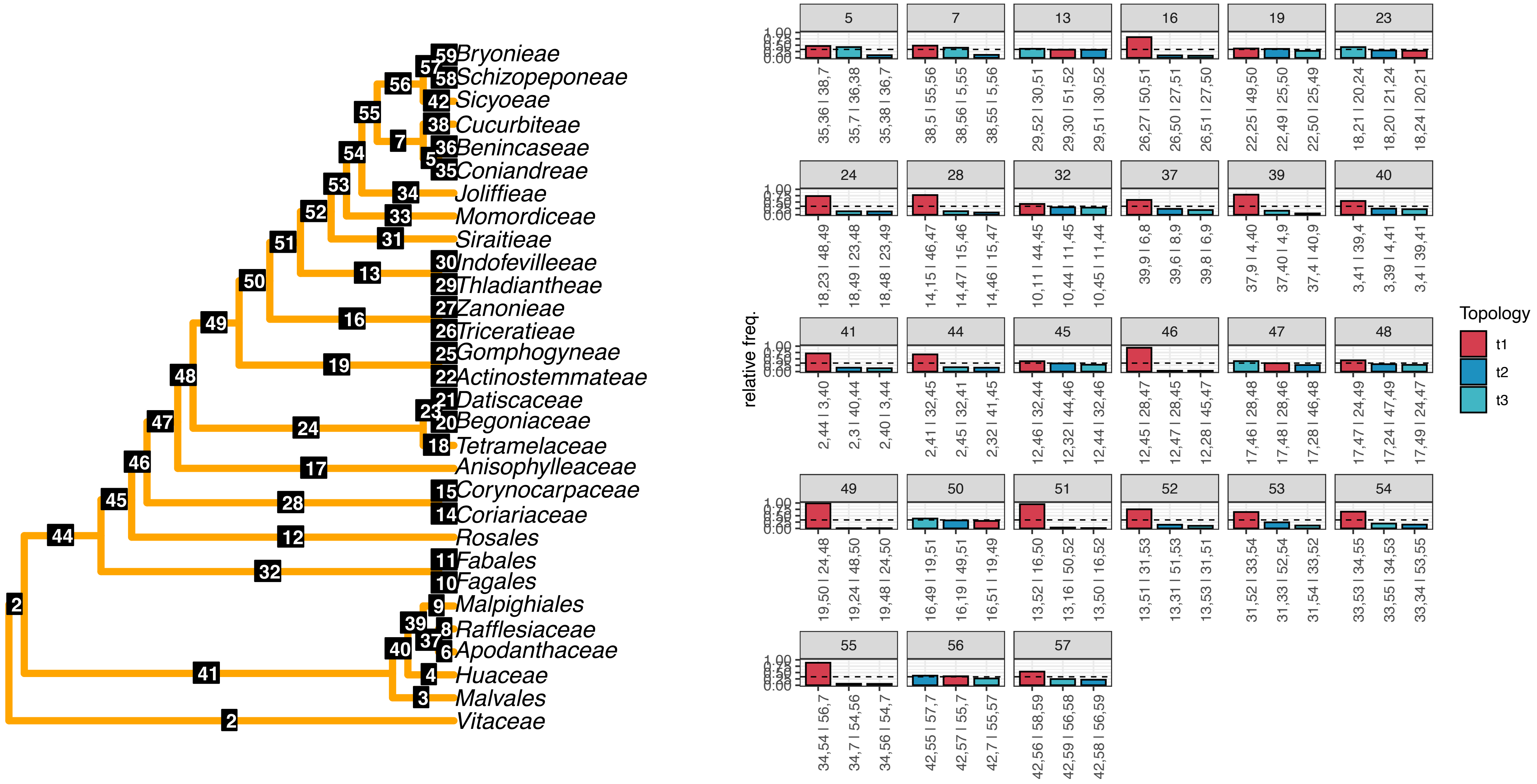

k) Method: FastTree
Dataset: Mega353
Paralog filter: informed

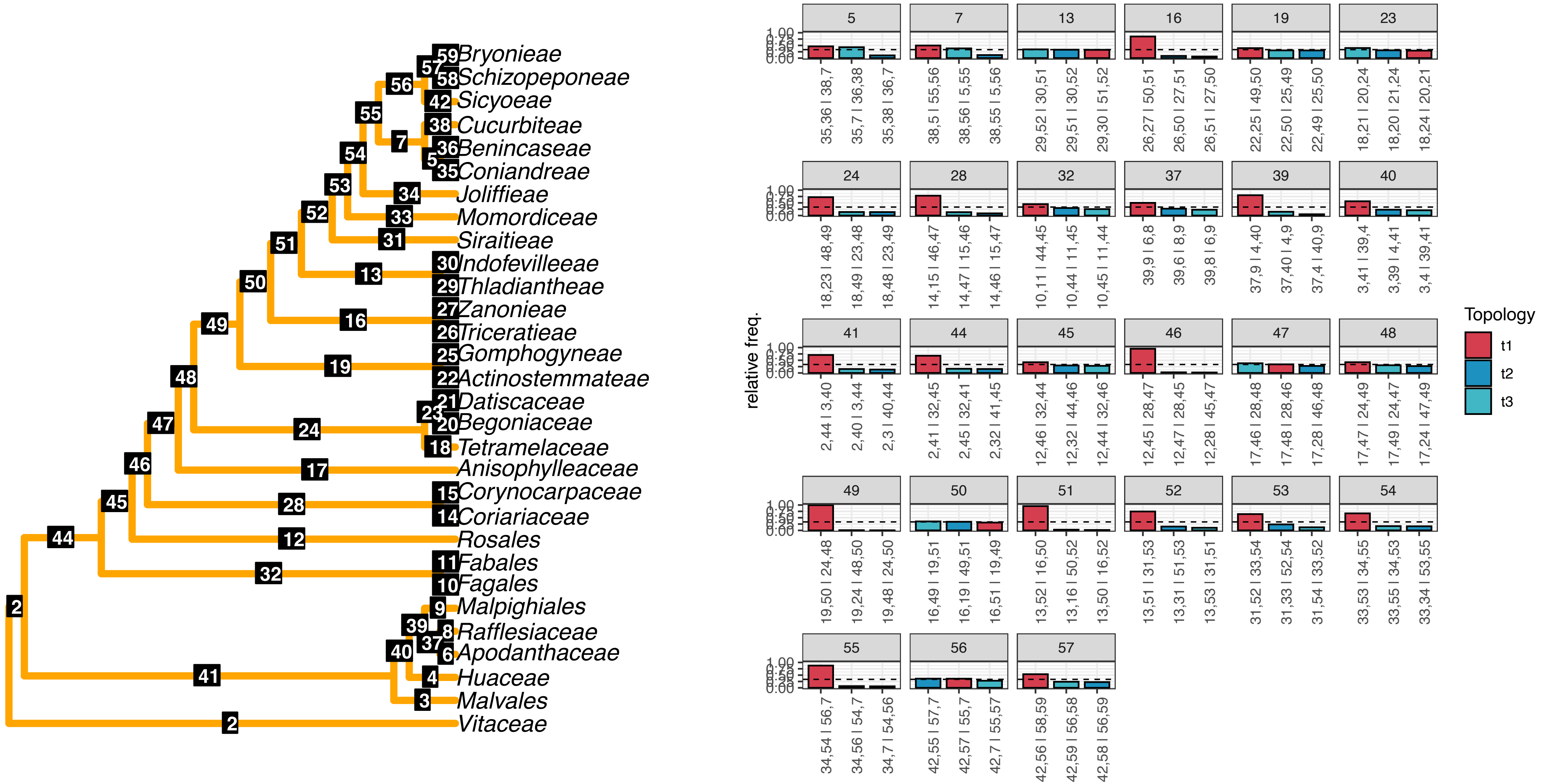

l) Method: FastTree

Dataset: Mega353

Paralog filter: naive

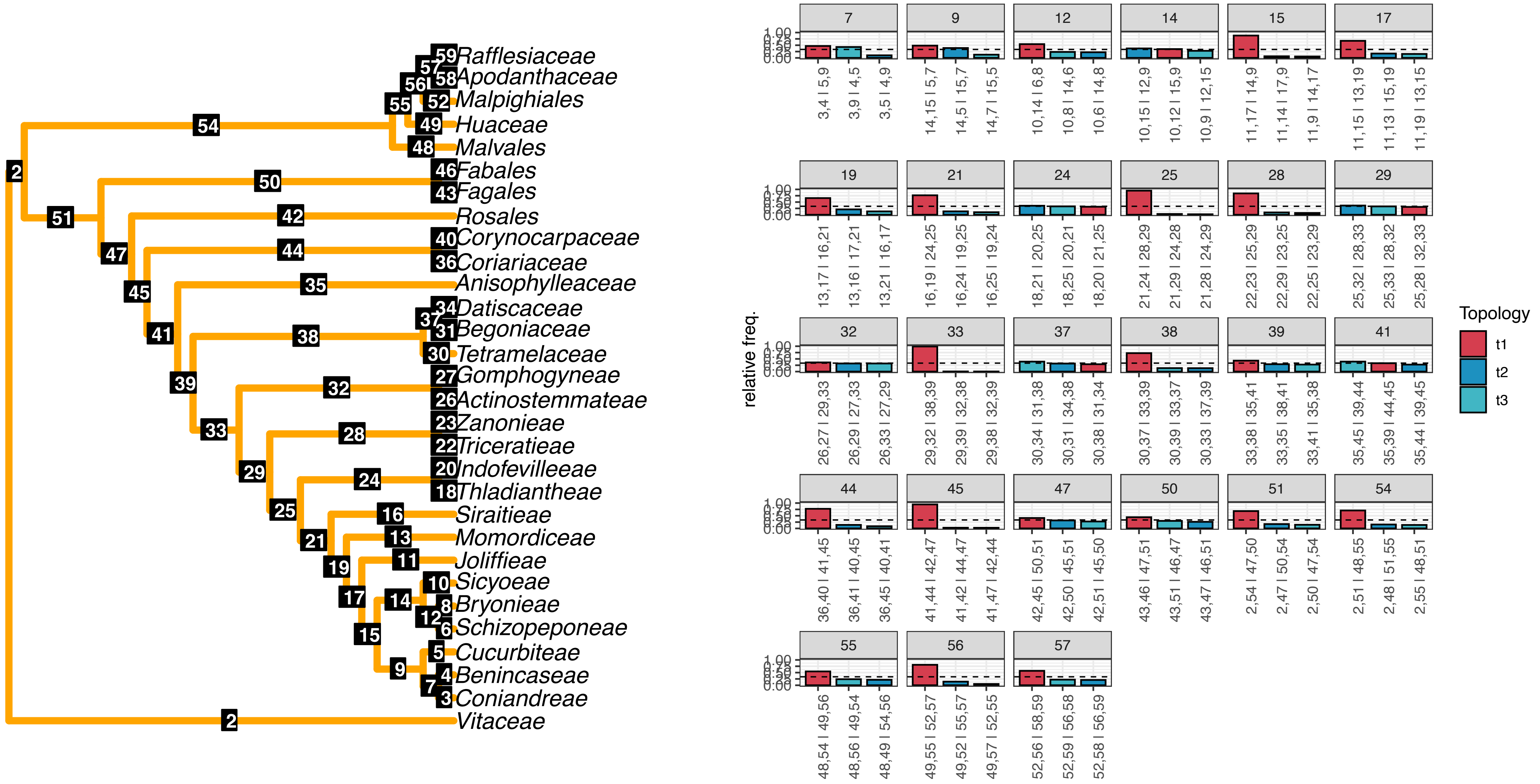
