## Supplementary material for "A novel phylogenomics pipeline reveals extensive topological conflict in the evolution of the angiosperm order Cucurbitales": SupplementaryMethod.pdf

**Supplementary Method:**

**Decontamination of the target capture sample *Lemurosicyos variegata***

The phylogenetic position of this sample in our preliminary analyses suggested contamination given that some gene trees were able to place it correctly inside its corresponding tribe Benincaseae in the Cucurbitaceae family while others place it outside the order Cucurbitales. Additionally, the extraction report from CAPTUS revealed that almost every gene had multiple copies recovered, suggesting a potential mixture of DNAs.

Since relatives in the tribe Benincaseae count with excellent genomic resources (e.g., *Cucumis sativus*) we can use one of these annotated genomes as a reference to retain only the contigs in the contaminated *Lemurosicyos* sample that contain high-identity hits to the nuclear and organellar proteins of *C. sativus*.

We first performed a CAPTUS extraction using as reference targets the nuclear proteins of *C. sativus* (Appendix S6), retaining at most two copies per recovered gene in case that both alleles were assembled correctly as separate contigs (`--max_paralogs 1`) and increasing the identity threshold to 85% (`--nuc_min_identity 85`) with the following command:

```
captus_assembly-runner.py extract \
      -a 02_assemble_Lemurosicyos_variegata \
      -o 03_extract_Lemurosicyos_variegata \
      -n ../references/cucumis_sativus/Cucumber_V3_chr_201810.pep.fa \
      --nuc_min_identity 85 \
      --max_paralogs 1 \
      --keep_all
```

Then we performed an additional extraction with CAPTUS targeting the organellar proteins of *C. sativus* as well as ribosomal genes and other non-coding regions from Cucurbitales species (Appendix S6), with the same increased identity but this time not allowing extra copies (`--max_paralogs 0`) with the following command:

```
captus_assembly-runner.py extract \
      -a 02_assemble_Lemurosicyos_variegata \
      -o 03_extract_Lemurosicyos_variegata \
      -p ../references/cucumis_sativus/Cucumis_sativus_PTD.faa \
```

```
        --ptd_min_identity 85 \
        -m ../references/cucumis_sativus/Cucumis_sativus_MIT.faa \
        --mit_min_identity 85 \
        -d ../references/Cucurbitales_NC.fna \
        --dna_min_identity 85 \
        --max_paralogs 0 \
        --keep_all \
        --overwrite
```

Finally, we used **only** the contigs that received hits during both extractions. After the procedure, the sample *Lemurosicyos variegata* is correctly and consistently placed as sister to an additional sample of the species which we sequenced at high-depth (Table S1, Table S4), both placed in the tribe Benincaseae.
