## Supplementary figures and images for "A novel phylogenomics pipeline reveals extensive topological conflict in the evolution of the angiosperm order Cucurbitales"

### FigS1.pdf

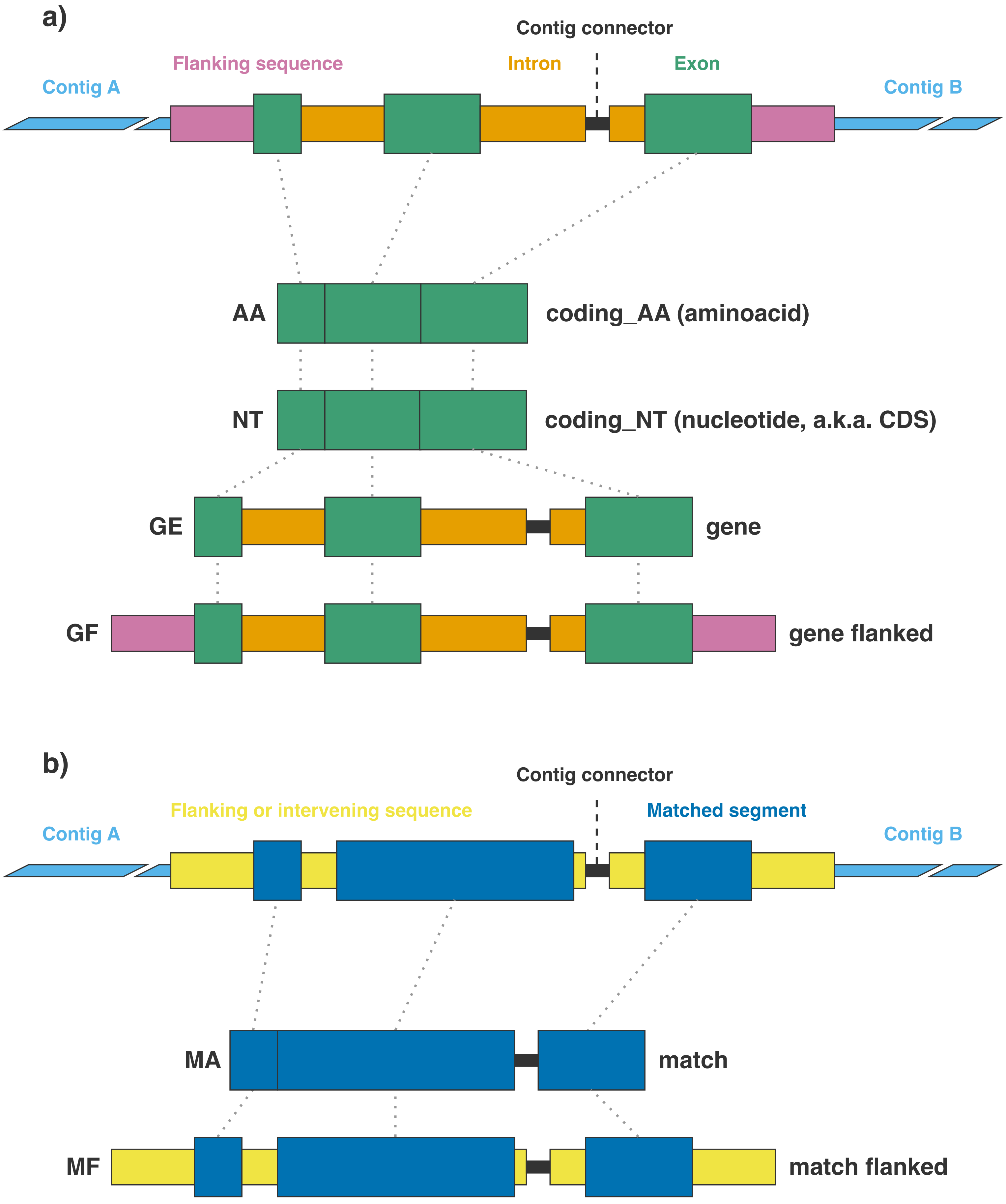

### FigS2.pdf

#### a) Dataset: Angiosperms353

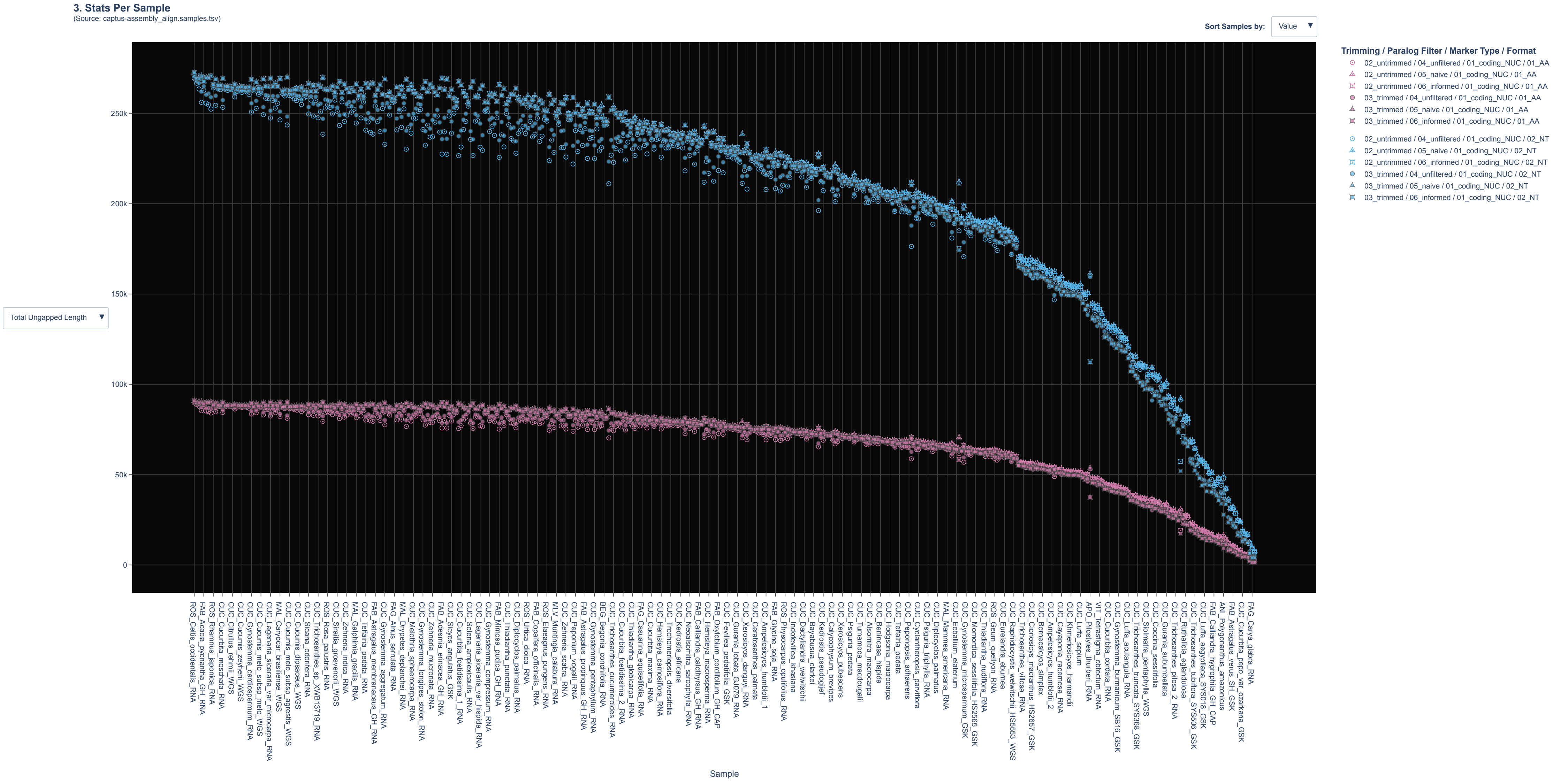

b) Dataset: Mega353

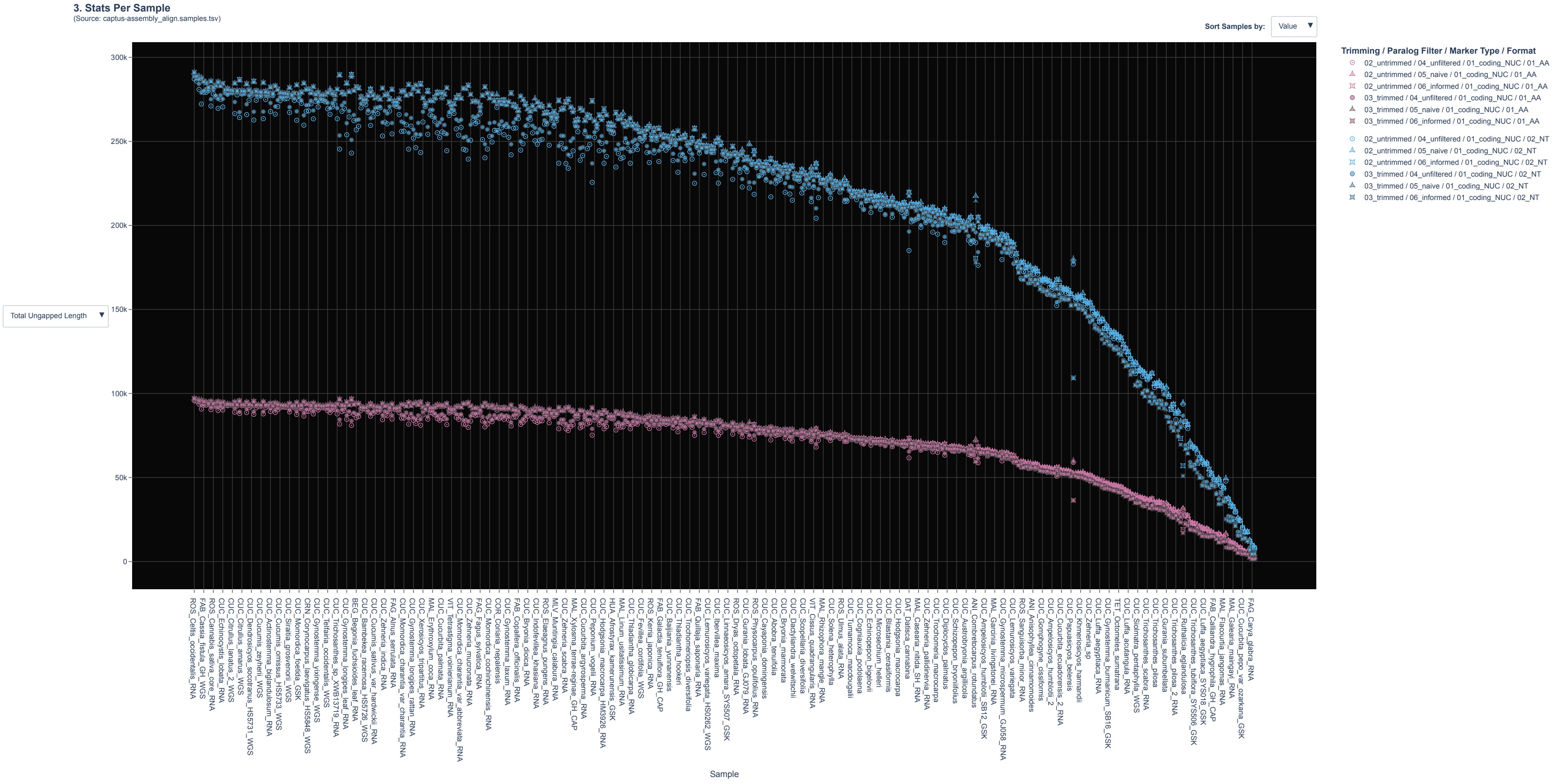

#### c) Dataset: RNA5435

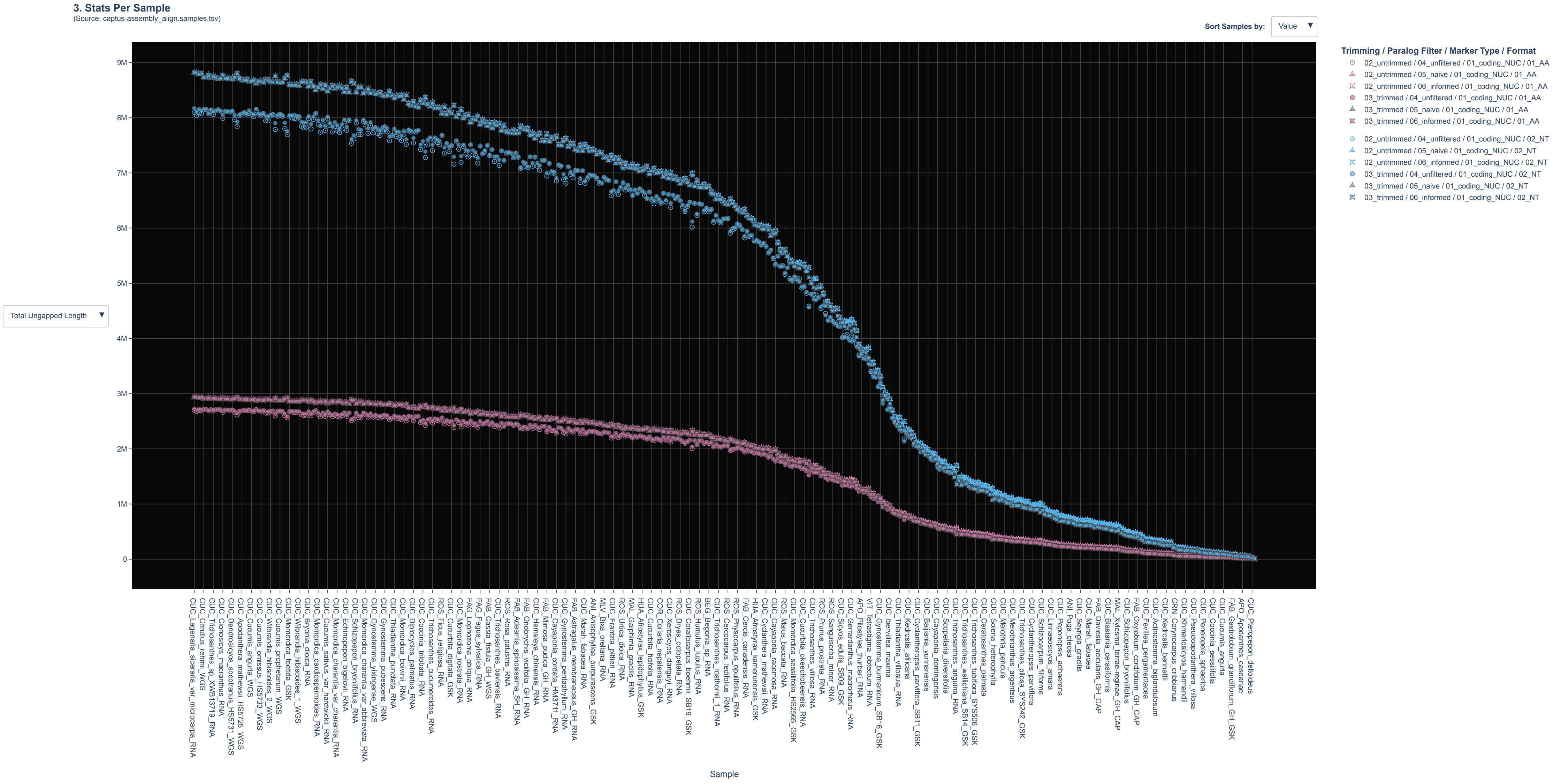

### FigS4.pdf

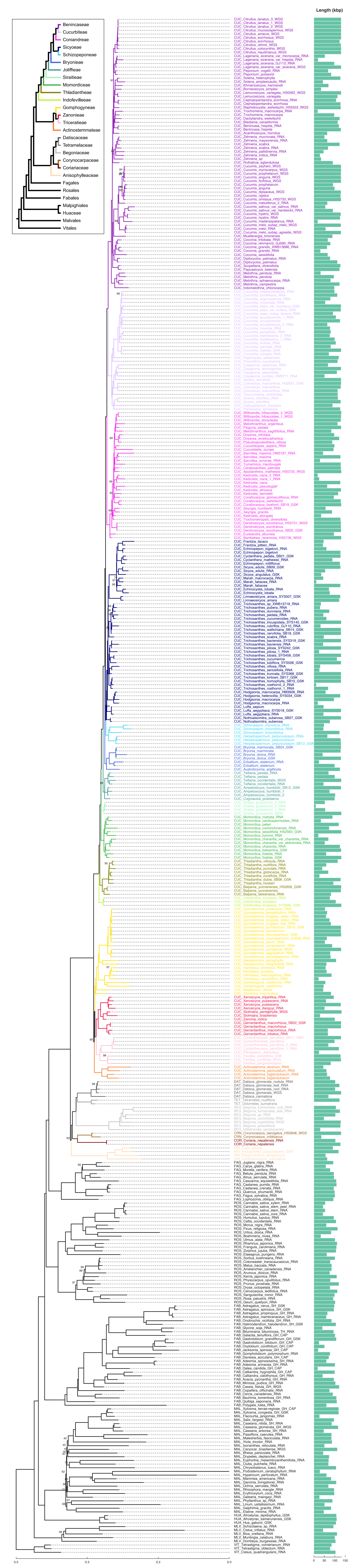

### FigS6.pdf

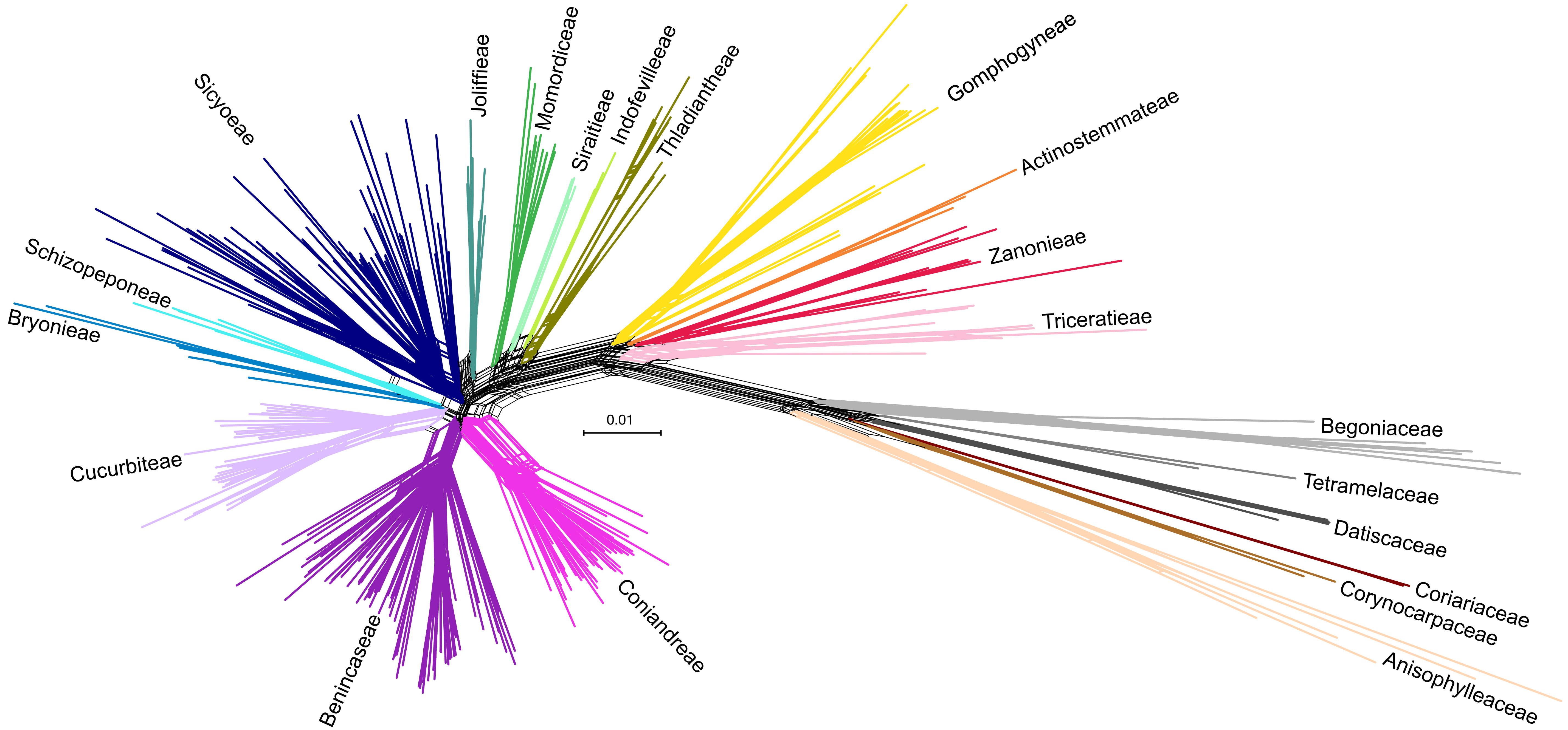

### FigS7.pdf

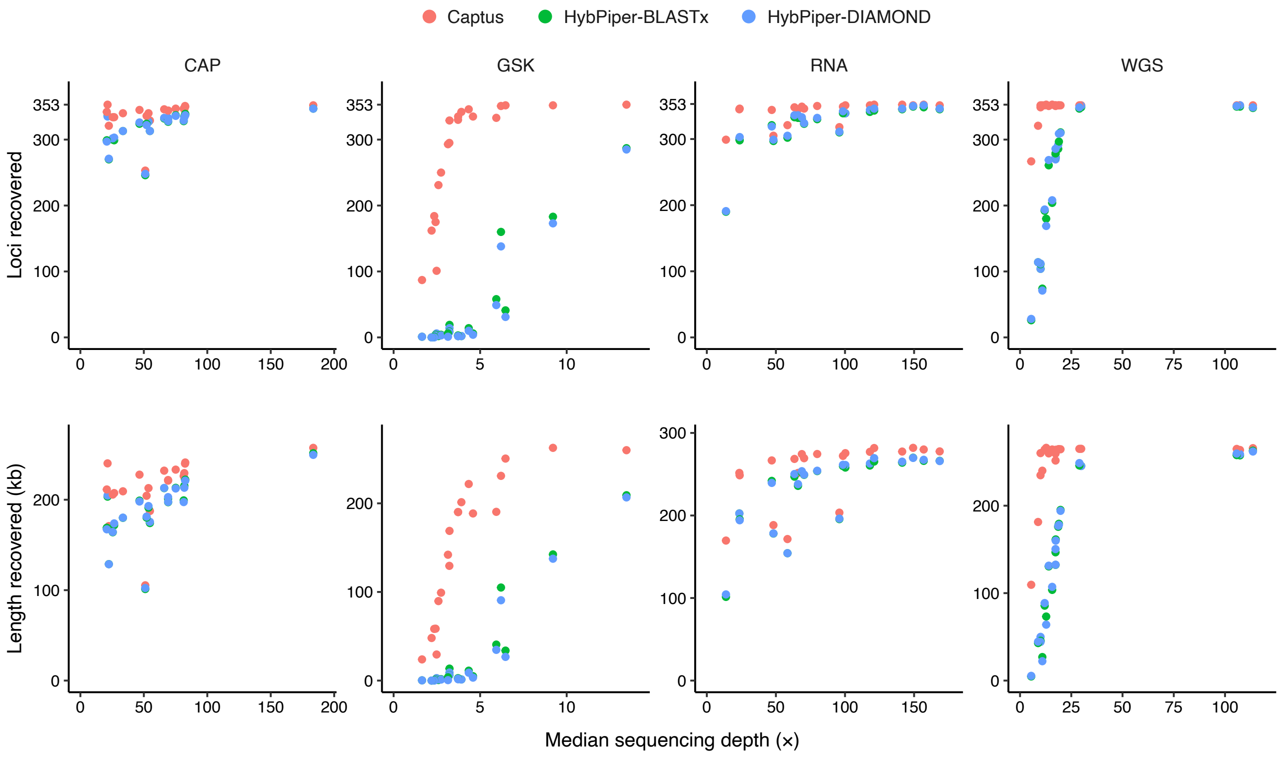

### FigS8.pdf

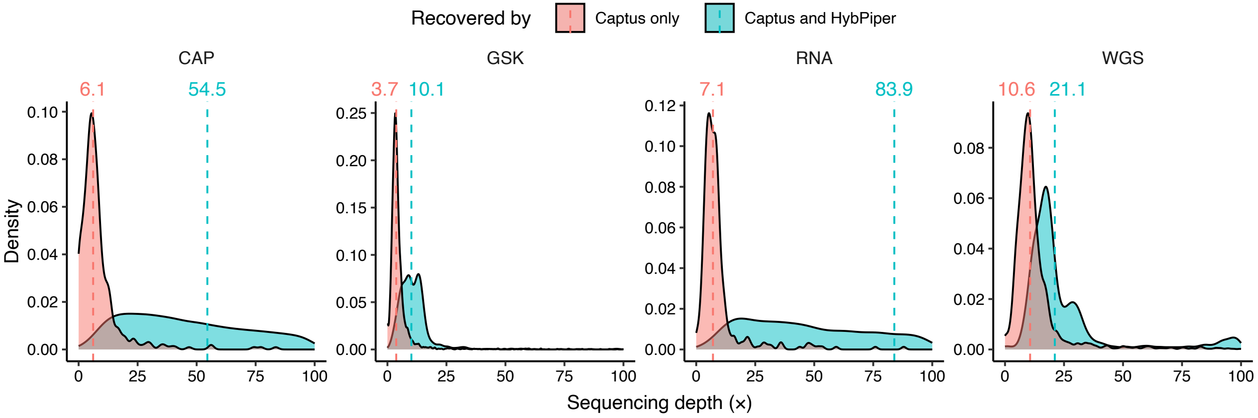

### FigS9.pdf

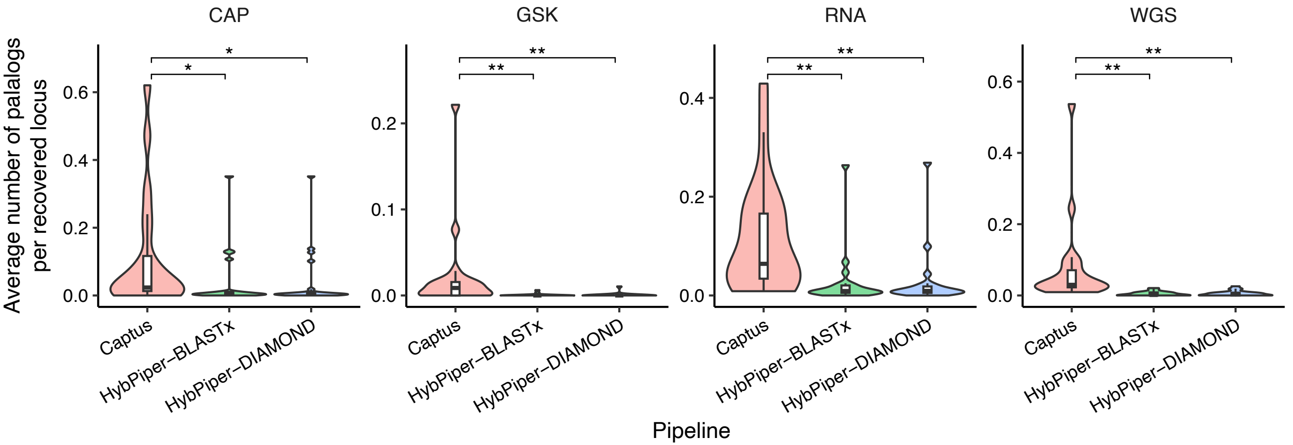

### FigS10.pdf

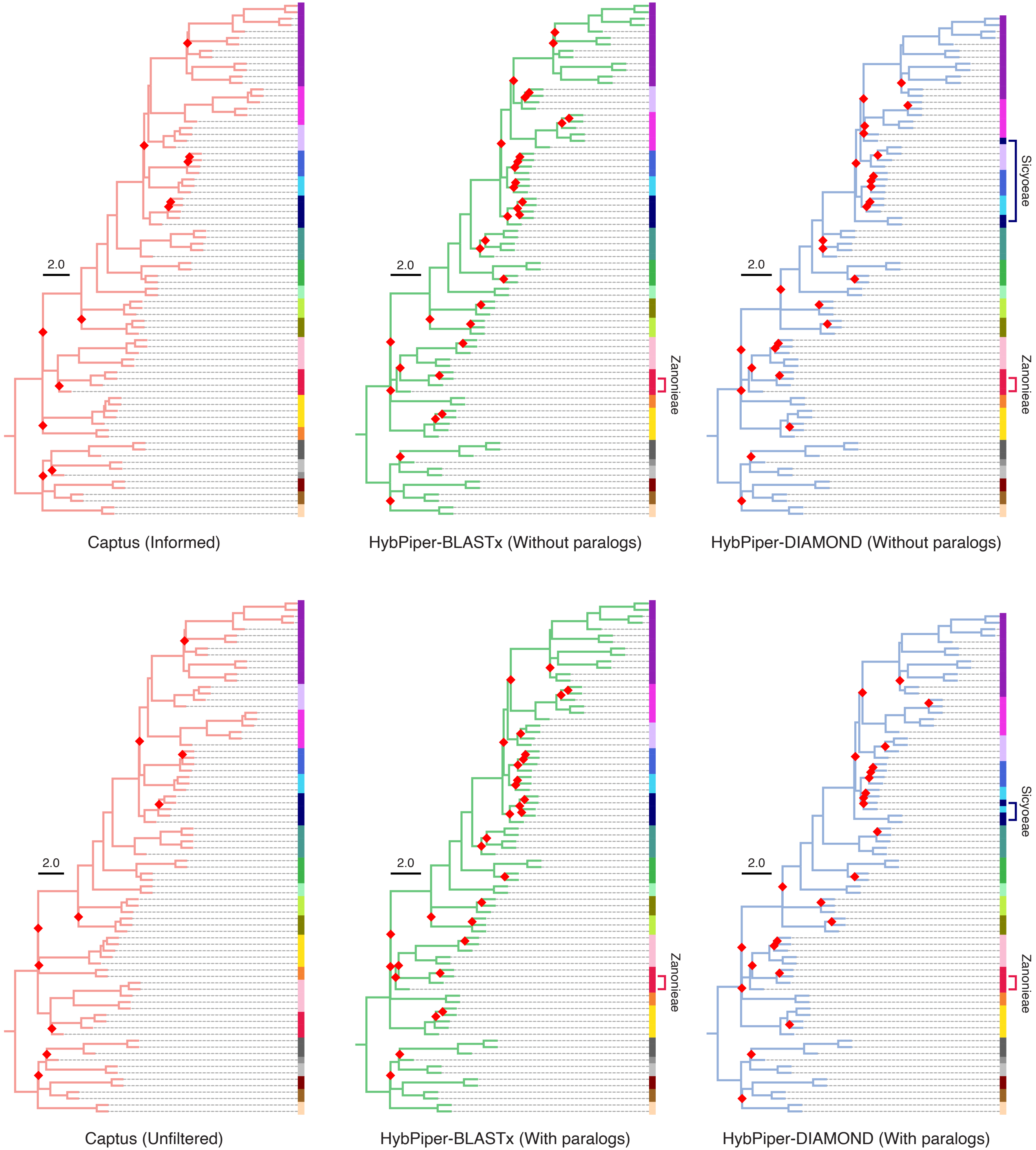
